# Targeting RAD52 overcomes PARP inhibitor resistance in preclinical *Brca2*-deficient ovarian cancer model

**DOI:** 10.1101/2025.09.24.678351

**Authors:** Yukihide Ota, Vijayalaxmi Gupta, Bisiayo E. Fashemi, Moreniola Akande, Preedia Babu, Prasanth Thuthika, Maia L. Elizagaray, Lulu Sun, Brooke Sanders, Lindsay M Kuroki, Carolyn K McCourt, Andrea R Hagemann, Ian S Hagemann, Premal H Thaker, David G Mutch, Matthew A Powell, Krzysztof Hyrc, Priyanka Verma, John Krais, Benjamin G. Bitler, Mary Mullen, Dineo Khabele

## Abstract

*BRCA*-mutated ovarian cancer commonly develops resistance to poly (ADP-ribose) polymerase (PARP) inhibitors. Here, we investigated the DNA repair protein RAD52 as a potential target to overcome resistance. In analysis of The Cancer Genome Atlas datasets and immunohistochemistry of tissue microarrays, elevated RAD52 expression correlated with poor overall survival in patients with high-grade serous ovarian cancers. We tested two PARP inhibitor-resistant *Brca2*-deficient mouse ovarian cancer models, ID8-OR and HGS2-OR. HGS2- OR cells had higher RAD52 expression than parental lines. *Rad52* knockout or knockdown restored PARP inhibitor sensitivity in both models. In syngeneic mice, ID8-OR cells in which *Rad52* was knocked out yielded lower tumor burden and longer overall survival than control cells. *Rad52* depletion impaired single-strand annealing and homologous recombination and led to accumulation of DNA double-strand breaks after PARP inhibitor treatment. RNA sequencing demonstrated that PARP inhibitor treatment induced *Polq* expression in *Brca2*- and *Rad52*-deficient cells, suggesting a switch to microhomology-mediated end joining. Finally, the RAD52 inhibitor D-I03 synergized with a PARP inhibitor to reduce cell viability and tumor burden and prolong survival. Collectively, our findings establish RAD52 as a promising therapeutic target to overcome PARP inhibitor resistance in *BRCA2*-mutated ovarian cancer and offer mechanistic insights to inform future clinical strategies.

## Introduction

Ovarian cancer is the sixth leading cause of cancer-related mortality among women in the United States, and approximately 20,890 new cases and 12,730 deaths are projected for 2025^1^. High-grade serous ovarian carcinoma (HGSC) is the predominant histologic subtype of epithelial ovarian cancer, accounting for nearly 70% of cases^2-4^. Approximately 50% of HGSC tumors exhibit homologous recombination (HR) repair deficiency (HRD), arising from diverse genetic and epigenetic alterations^2^. Germline or somatic mutations in *BRCA1* or *BRCA2* are observed in ∼10% of cases, and epigenetic silencing of *BRCA1* via promoter hypermethylation accounts for an additional ∼10%. The remaining HRD cases are attributed to defects in other HR pathway genes, including *RAD51C*, *RAD51D*, and *PALB2*^2^.

The current standard treatment for HGSC is cytoreductive surgery and platinum-based chemotherapy, which initially yields high response rates^5,6^. Additionally, poly (ADP-ribose) polymerase inhibitors (PARPis) are used as maintenance therapy or as treatment for recurrent disease. PARPis have shown substantial efficacy in patients whose tumors have *BRCA1/2* mutations or exhibit HRD^7-9^. For example, in newly diagnosed advanced ovarian cancer with *BRCA1/2* mutations, maintenance olaparib extended progression-free survival by 42.2 months^10^. Despite these benefits, resistance to PARPis is a major clinical challenge, with approximately 50% of *BRCA*-mutated ovarian cancer patients relapsing within five years of initiating PARPi therapy^10^. Notably, there is currently no approved targeted therapy specifically designed to overcome acquired PARPi resistance in *BRCA*-mutated ovarian cancer.

Mechanisms of PARPi resistance in HRD ovarian cancer can be broadly classified into two major categories^11,12^. The first involves restoration of HR function, commonly through *BRCA1/2* reversion mutations^13-15^, alternative splicing, hypomorphic BRCA1 activity^16,17^, or loss of *BRCA1* promoter hypermethylation^18^. The second category includes HR-independent mechanisms such as increased expression of drug efflux transporters (e.g., ABCB1)^19,20^, stabilization of replication forks, or suppression of replication-coupled gaps^21^. HR-deficient tumors that develop PARPi resistance via HR-independent mechanisms may necessitate different therapeutic strategies than tumors with restored or inherent HR proficiency^22-27^. Targets proposed to overcome PARPi resistance include DNA polymerase theta (gene *POLQ*)^28-31^ and ATR^32,33^, whose protein overexpression promotes microhomology-mediated end joining (MMEJ) and the replication stress response, respectively. Given the heterogeneity of drug resistance mechanisms among patients and cell types, personalized therapeutic approaches are critically needed.

An alternative DNA repair pathway that shows potential as a target in HR-deficient cells is regulated by RAD52, since targeting of RAD52 is synthetically lethal with loss of BRCA2^34^. RAD52 promotes single-strand annealing (SSA) through its N-terminal domain, which binds single-stranded DNA (ssDNA) and facilitates error-prone repair by using homologous sequences of more than 29 base pairs^35,36^. The N-terminal domain is also responsible for oligomerization, forming ring structures that are associated with SSA activity^35-37^. This domain also supports break-induced replication, mitotic DNA synthesis, RNA-mediated repair of DNA, and replication fork protection under replication stress^38-42^. The C-terminal domain of RAD52 contains binding motifs for RAD51 and replication protein A and a nuclear localization signal. In yeast, RAD52 promotes RAD51 filament formation during HR, but its role in mammals is unclear, especially given the poor conservation of the RAD51-binding domain^43^. Pharmacologic inhibition of RAD52 enhances the efficacy of PARPis in *BRCA*-deficient cells^44-46^. However, whether targeting RAD52 enhances the efficacy of PARPis in acquired PARPi-resistant cells and the specific contribution of RAD52 to PARPi resistance in *BRCA2*-mutated ovarian cancer is unknown.

In this study, we used public databases and immunohistochemistry of a tissue microarray to show that high RAD52 expression correlates with poor prognosis in ovarian cancer patients. Depletion or inhibition of RAD52 re-sensitized mouse *Brca2*-deficient, PARPi-resistant ovarian cancer cells to PARPis both *in vitro* and *in vivo*. *Rad52* depletion reduced both SSA and HR activity and led to accumulation of DNA damage after PARPi treatment. In the HGS2-OR cell line, elevated RAD52 expression was associated with increased SSA and HR activity, suggesting that RAD52 upregulation partially contributes to PARPi resistance. Furthermore, RNA-seq analysis revealed upregulation of *Polq* in *Rad52* knockout cells treated with PARPis, suggesting that the MMEJ pathway is activated as a compensatory DNA repair mechanism in the absence of *Rad52*. Together, our findings reveal RAD52 as a promising therapeutic target for overcoming PARPi resistance in *BRCA2*-deficient ovarian cancer.

## Results

### High RAD52 expression is associated with poor overall survival in patients with high-grade serous ovarian carcinoma (HGSC)

We investigated the association between RAD52 protein expression and overall survival in patients with high-grade serous ovarian carcinoma (HGSC) by immunohistochemistry of a tissue microarray comprising HGSC samples (Supplementary Fig. 1, Supplementary Table 1). Patients were stratified according to the RAD52 H-score, using the median value of 168 as the cutoff (Fig. 1a). In Kaplan–Meier survival analysis, high RAD52 protein expression (n = 36) was associated with significantly worse 4-year overall survival compared with low RAD52 protein expression (n = 36) (58.3% vs. 80.6%, log-rank p = 0.03), although the difference was not significant at 5 years (Fig. 1b). To validate these findings in a larger cohort, we analyzed data from The Cancer Genome Atlas (TCGA). We included patients with ovarian serous cystadenocarcinoma harboring *TP53* mutations, as 96% of HGSC tumors carry such alterations^47^. Patients were stratified into high and low *RAD52* expression groups based on median mRNA levels obtained from RNA-seq data. High *RAD52* expression was significantly associated with shorter overall survival compared with low expression (median 36.3 vs. 49.5 months, log-rank p = 0.02) (Fig. 1a). Collectively, these results demonstrate that elevated RAD52 expression correlates with poor prognosis in HGSC.

**Figure 1.**
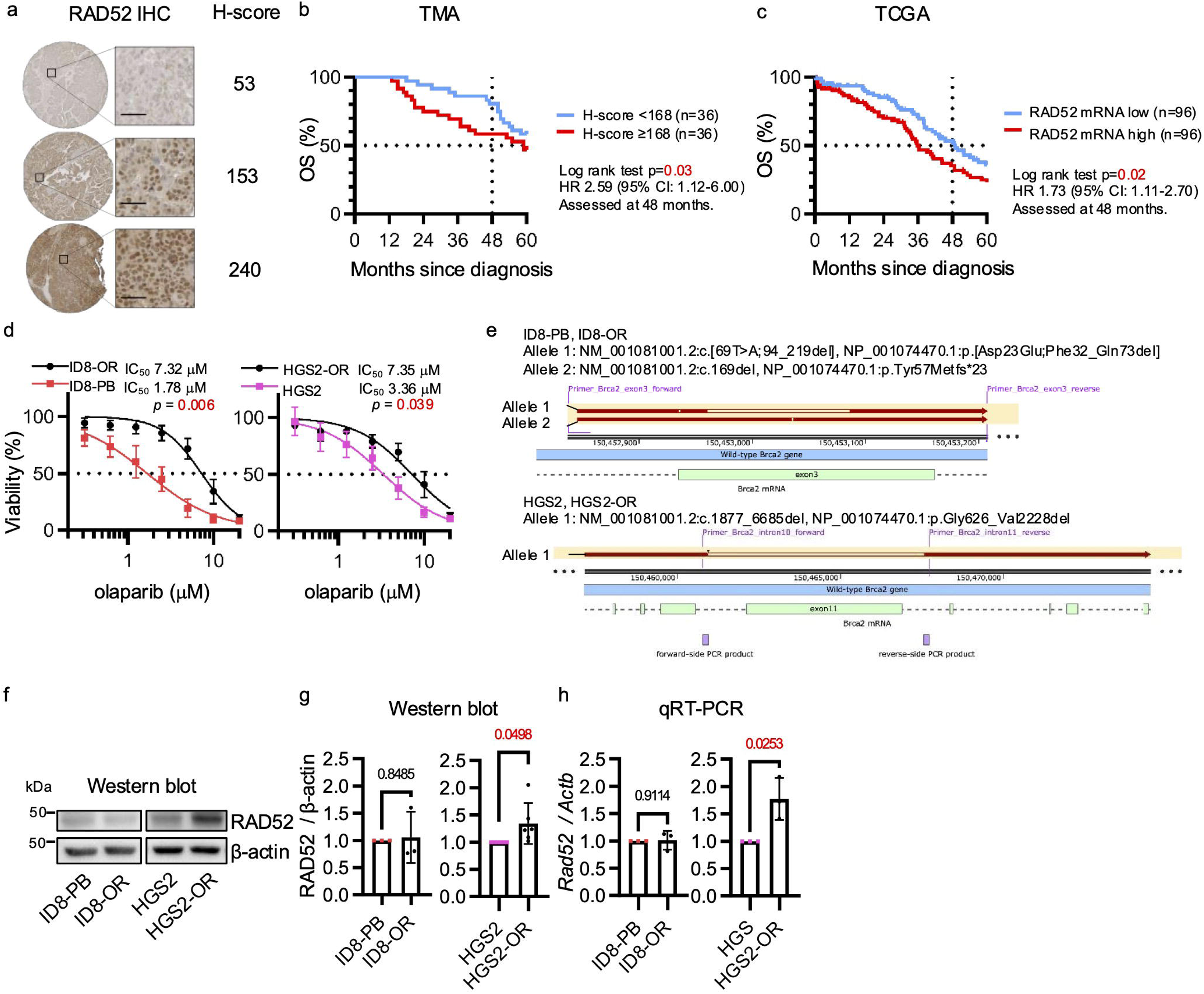
RAD52 is associated with poor prognosis in serous ovarian cancer and is upregulated in a *BRCA2*-deficient, PARP inhibitor–resistant cell line. **(a)** Representative immunohistochemical images of RAD52 staining in the tissue microarray. **(b)** Kaplan-Meier analysis of OS of patients with high grade serous ovarian cancer whose tumors were represented in the tissue microarray dichotomized as high (H-score ≥ 168) or low (H-score < 168) RAD52 protein expression. The log-rank test was used to analyze survival at 4 years after diagnosis. **(c)** Kaplan-Meier analysis of overall survival (OS) of patients from the The Cancer Genome Atlas database with serous ovarian cancer with *TP53* mutations and high or low *RAD52* mRNA expression. The log-rank test was used to analyzed survival at 4 years after diagnosis. **(d)** Viability assays comparing *Brca2*-deficient parental (PB) and olaparib-resistant (OR) ovarian cancer cells treated with olaparib for 5 days. **(e)** *Brca2* mutation status was evaluated by next-generation sequencing. Exon 3 was analyzed in ID8-PB and ID8-OR cells, and exon 11 in HGS2 and HGS2-OR cells. (f) Western blots of RAD52 protein expression in *Brca2*-deficient parental and OR cells. **(g, h)** Quantification of RAD52 **(g)** protein and **(h)** mRNA normalized to β-actin and shown relative to the amount in parental cells. Data show mean ± standard deviation of at least three independent experiments, with individual dots for each replicate. Statistical significance was assessed by two-sided Student’s t-tests.

### RAD52 is upregulated in a *Brca2*-deficient, PARP inhibitor–resistant mouse ovarian cancer cell line

We used two olaparib-resistant mouse ovarian cancer cell lines, ID8-OR and HGS2-OR, which were previously generated by exposing the parental cell lines ID8-PB (*Trp53*^−/−^, *Brca2*^−/−^) and HGS2 (*Trp53*^−/−^, *Brca2*^−/−^, *Pten*^−/−^) to olaparib and were characterized by Benjamin G. Bitler et al^48,49^. Both OR cell lines showed a significantly higher olaparib IC₅₀ than their parental counterparts (Fig. 1d), indicating acquired resistance. To rule out *Brca2* reversion mutations as a resistance mechanism, we performed next-generation sequencing. This confirmed that the ID8-OR cells carried the same deletion or frameshift mutation in exon 3 of the *Brca2* gene as the parental ID8-PB cells (Fig. 1e, Supplementary Fig. 2). Similarly, the HGS2-OR cells had the same over 4-kb deletion within exon 11 as the parental HGS2 cells. Exon 3 of BRCA2 encodes the PALB2-binding domain, whereas exon 11 contains the BRC repeats required for RAD51 binding, both of which are critical for homologous recombination^50,51^. Therefore, these results confirm that PARPi resistance in these cell lines was not driven by *Brca2* reversion mutations. Next, RAD52 protein and mRNA were quantified by Western blotting and quantitative reverse transcription PCR (qRT-PCR), respectively, and all values were normalized to the corresponding parental values. HGS2-OR cells exhibited higher RAD52 protein (fold change ∼1.34) and mRNA expression (fold change ∼1.77) than their corresponding parental cells, whereas ID8-OR cells did not show such upregulation (Fig. 1f-g). High RAD52 expression was thus a feature of one PARPi-resistant *Brca2*-deficient ovarian cancer cell line.

### Genetic depletion of *Rad52* restores PARPi sensitivity in *Brca2*-deficient PARPi-resistant ovarian cancer cell lines

To investigate whether genetic depletion of *Rad52* restored olaparib sensitivity in *Brca2*-deficient, PARPi–resistant ovarian cancer cells, we used CRISPR/Cas9 to generate two independent *Rad52* knockout (KO) clones (KO#1, KO#2) in ID8-OR cells. Additionally, we used short hairpin RNA (shRNA) to establish a stable knockdown (KD) clone (KD#1) in HGS2-OR cells. The structure of *Rad52* and the truncated *Rad52* resulting from knockout are shown in Supplementary Fig. 3. The efficiency of *Rad52* depletion was confirmed by Western blotting, qRT-PCR, and amplicon sequencing to verify genomic DNA editing (Fig. 2a, 2b, Supplementary Fig. 3). *Rad52*-depleted cells had significantly lower olaparib IC₅₀ (Fig. 2c) and formed significantly fewer clones than Scr controls upon olaparib treatment (Fig. 2d, 2e). These findings indicate that genetic depletion of *Rad52* re-sensitizes PARPi-resistant *Brca2*-deficient ovarian cancer cells to PARPi.

**Figure 2.**
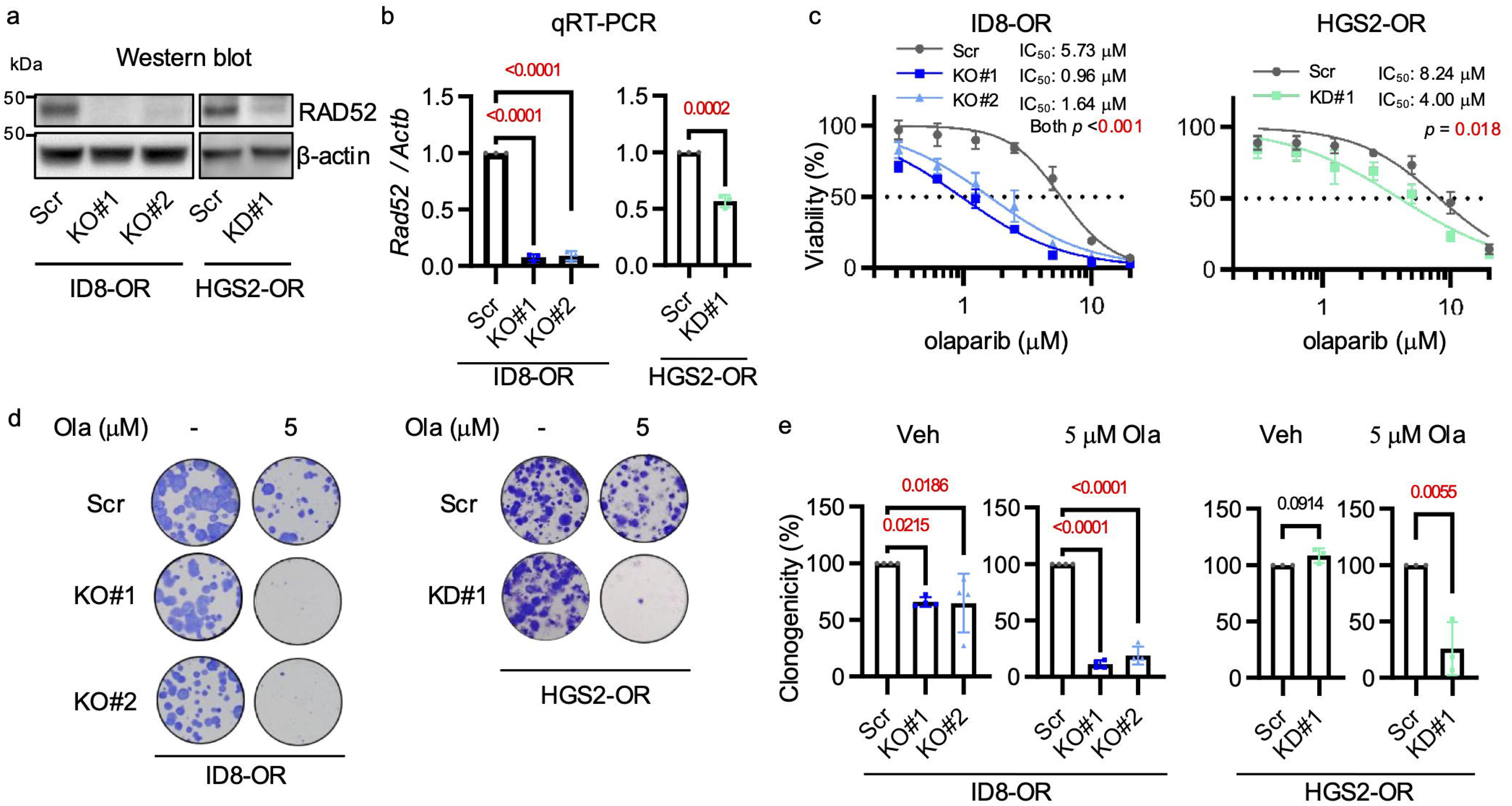
Genetic depletion of *Rad52* restores PARP inhibitor sensitivity in *Brca2*-deficient, PARP inhibitor–resistant ovarian cancer cells. **(a)** Western blot validation of *Rad52* knockout in ID8-OR cells and knockdown in HGS2-OR cells. **(b)** qRT-PCR of *Rad52* mRNA normalized to *Actb*. One of the primers was designed to span the CRISPR/Cas9 cleavage site. **(c)** Cell viability assay comparing *Rad52*-depleted cells with Scr control cells treated with a range of 0–20 µM olaparib for 5 days. **(d)** Clonogenicity assay following 8-day treatment with 5 μM olaparib or vehicle. **(e)** Quantification of clonogenicity assay from panel (**d**). Data show mean ± standard deviation of at least three independent experiments, with individual dots for each replicate. Statistical significance was assessed by two-sided Student’s t-tests for two-group comparisons and one-way ANOVA followed by Šídák’s multiple comparisons test for comparisons among three groups. Veh, vehicle; Ola, olaparib.

### Genetic depletion of *Rad52* reduces tumor burden and ascites in a *Brca2*-deficient, PARP inhibitor–resistant ovarian cancer model

We evaluated the impact of genetic depletion of *Rad52* on tumor progression in a mouse model of *Brca2*-deficient, PARPi–resistant ovarian cancer. ID8-Scr or ID8-KO#1 cells (5 × 10⁶) were intraperitoneally injected into C57BL/6 mice. Beginning one week after tumor cell inoculation, olaparib (50 mg/kg) or vehicle were administered daily for four weeks (Fig. 3a), and then tumor burden and ascites volume were assessed. In both vehicle- and olaparib-treated groups, the ID8-KO#1 group showed significantly lower total tumor weight, omental weight, number of nodules, and ascites volume than the Scr group (Fig. 3b-f, Supplement Fig. 4). Olaparib-treated *Rad52* KO#1 tumors had a significantly higher percentage of cells that were positive for the DNA double-strand break marker γH2AX^52,53^ than did their untreated counterparts. Furthermore, the proportion of γH2AX-positive cells was higher in *Rad52* KO#1 tumors than in Scr tumors, regardless of olaparib treatment (Fig. 3g, 3h). Kaplan–Meier survival analysis revealed that mice receiving *Rad52*-KO#1 cells had longer overall survival than those receiving Scr control cells (Fig. 3i). These results indicate that loss of *Rad52* suppresses tumor burden and improves survival in a mouse model of *Brca2*-deficient, PARPi–resistant ovarian cancer.

**Figure 3.**
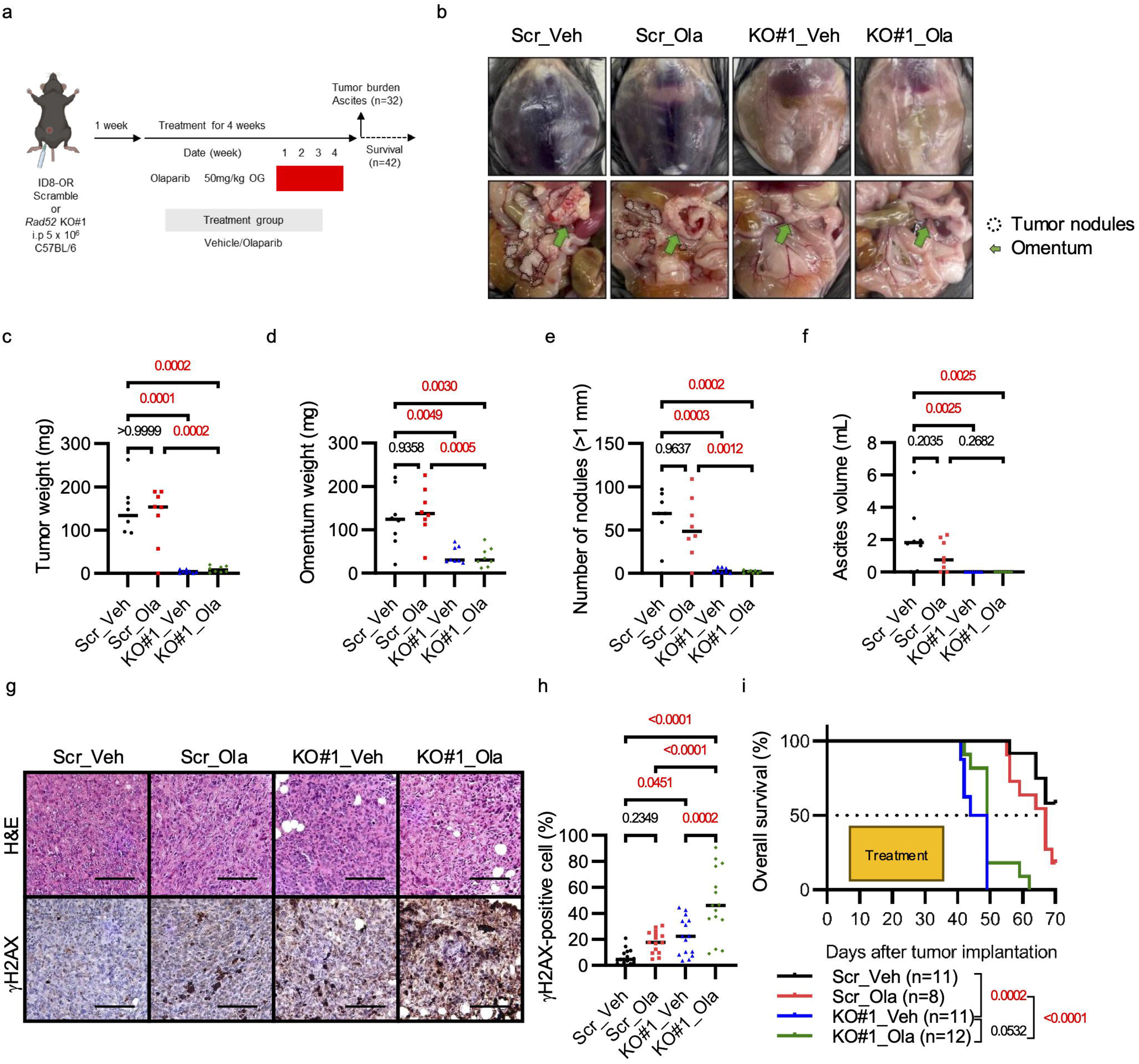
*Rad52* depletion suppresses tumor growth in a mouse model of *Brca2*-deficient ovarian cancer. **(a)** Schematic illustrating the ID8-OR xenograft mouse model experimental design. **(b)** Representative macroscopic images of mice in the four groups. The dotted areas indicate disseminated nodules in the peritoneal cavity. Green arrows indicate the omentum. **(c)** Total tumor weight excluding the omentum. Mean ± standard deviation: Scr_Veh 132.4 ± 76.1 mg, Scr_Ola 131.6 ± 68.1 mg, KO#1_Veh 4.4 ± 4.0 mg, KO#1_Ola 9.5 ± 7.0 mg. **(d)** Weight of the omentum (including the pancreas and surrounding adipose tissue). Scr_Veh 125.0 ± 67.1 mg, Scr-Ola 141.0 ± 57.1 mg, KO#1-Veh 41.5 ± 19.5 mg, KO#1-Ola 37.0 ± 22.3 mg. **(e)** Number of intraperitoneal tumor nodules (≥1 mm in diameter). Scr_Veh 60.3 ± 35.4, Scr_Ola 53.0 ± 34.7, KO#1_Veh 3.4 ± 2.9, KO#1_Ola 1.8 ± 1.6. **(f)** Volume of ascites. Scr_Veh 2.1 ± 2.0 mL, Scr_Ola 1.0 ± 0.9 mL, KO#1_Veh 0 ± 0 mL, KO#1_Ola 0 ± 0 mL. For panels (**c**)–(**f**), n = 8 mice per group. **(g)** Representative H&E and immunohistochemical staining for γH2AX. Scale bar, 100 μm. **(h)** Quantification of γH2AX -positive nuclei in intraperitoneal tumor tissues. P-values determined by one-way ANOVA followed by Šídák’s multiple comparisons test. **(i)** Kaplan–Meier survival curves. A log-rank test was initially conducted across all four groups, followed by pairwise comparisons between selected groups. To account for multiple comparisons (three in total), the significance threshold was adjusted to *p* < 0.0167 using Bonferroni correction (*0.05/3*). Median survival (days): Scr_Veh 49, Scr-Ola 46.5, KO#1-Veh 67, KO#1-Ola Not reached.

### Genetic depletion of *Rad52* decreases single-strand annealing and homologous recombination activity and increases accumulation of DNA damage in *Brca2*-deficient, PARP inhibitor–resistant ovarian cancer cell lines

To determine which DNA repair pathways are affected by Rad52 depletion and whether olaparib-resistant cells exhibit elevated SSA and HR activity, we performed reporter assays to evaluate the repair of DNA double-strand breaks. In the SSA reporter assay, cells were co-transfected with the hprtSA-GFP construct, which contains an I-SceI recognition site, and the pCBASceI plasmid, which expresses the I-SceI endonuclease (Fig. 4a, 4b). GFP expression is restored upon SSA-mediated repair of an I-SceI–induced double-strand break. All data were normalized to the corresponding OR or Scr control cells. In addition, ID8-P cells (*Trp53^⁻/⁻^*, *Brca2^⁺/⁺^*), an HR-proficient ovarian cancer model, were included as a positive control for HR, together with ID8-PB cells (*Trp53^⁻/⁻^*, *Brca2^⁻/⁻^)*. SSA activity was significantly lower in ID8-P cells than in ID8-OR cells, indicating a reduced dependency on SSA in HR-proficient ID8-P cells (Fig. 4c). Consistent with the comparable RAD52 expression between ID8-PB and ID8-OR cells (Fig. 1f–h), no significant difference in SSA activity was observed between these two cell lines (Fig. 4c). In contrast, HGS2-OR cells, which express higher levels of RAD52 than parental HGS2 cells (Fig. 1f–h), displayed significantly increased SSA activity (Fig. 4d). Furthermore, ID8-KO#1, ID8-KO#2, and HGS2-KD#1 cells exhibited significantly reduced SSA activity compared with ID8-Scr or HGS2-Scr cells (Fig. 4e, 4f, Supplementary Fig. 5). Next, we assessed HR activity using the pDR-GFP construct, following the same procedure as in the SSA reporter assay (Fig. 4g). HR activity was significantly higher in ID8-P cells than in ID8-OR cells, consistent with the presence of wild-type *Brca2* in ID8-P cells (Fig. 4h). No significant difference in HR activity was observed between ID8-PB and ID8-OR cells. In contrast, HGS2-OR cells exhibited significantly higher HR activity than parental HGS2 cells (Fig. 4i). Similarly, ID8-KO#1, ID8-KO#2, and HGS2-KD#1 cells showed significantly reduced HR activity compared with ID8-Scr or HGS2-Scr cells (Fig. 4j, 4k, Supplementary Fig. 5).

**Figure 4.**
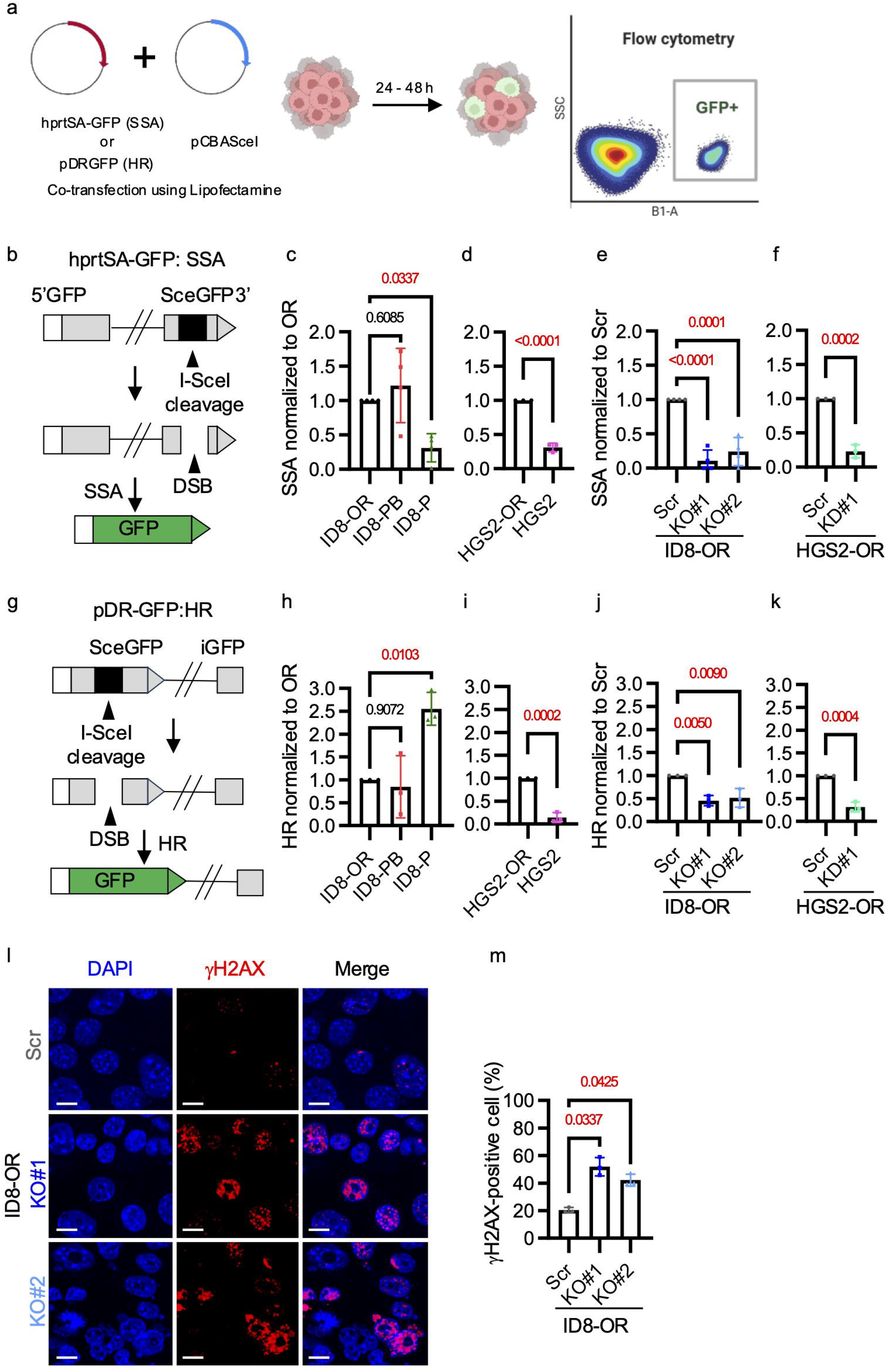
RAD52 promotes SSA and HR repair activity and limits DNA damage accumulation in PARP inhibitor-resistant cells. **(a)** Reporter plasmids used in GFP-based assays to measure single-strand annealing (SSA) and homologous recombination (HR) activity. **(a)** Schematic illustration of the GFP-based reporter assays. Reporter plasmids were co-transfected with pCBASceI, which encodes the I-SceI endonuclease. Successful repair of I-SceI–induced double-strand breaks restore GFP expression, which can be detected by flow cytometry 24–48 hours after transfection. **(b)** hprtSA-GFP construct used in GFP-based assays to measure single-strand annealing (SSA) **(c)** SSA activity in ID8-OR (*Trp53^⁻/⁻^*, *Brca2^⁻/⁻^*, olaparib-resistant), ID8-PB (*Trp53*^⁻/⁻^, *Brca2*^⁻/⁻^), and ID8-P (*Trp53*^⁻/⁻^, *Brca2*^⁺/⁺^) cells, normalized to ID8-OR**. (d)** SSA activity in HGS2-OR (*Trp53^⁻/⁻^*, *Brca2^⁻/⁻^*, *Pten*^-/-^, olaparib-resistant) and HGS2 (*Trp53*^⁻/⁻^, *Brca2*^⁻/⁻,^ *Pten*^-/-^) cells, normalized to HGS2-OR**. (e)** SSA activity in ID8-Scr, ID8-*Rad52*KO#1, and ID8-*Rad52*KO#2, normalized to ID8-Scr. **(f)** SSA activity in HGS2-Scr, HGS2-*Rad52*KD#1, normalized to HGS2-Scr. **(g)** pDR-GFP construct used in GFP-based assays to measure homologous recombination (HR). **(h)** HR activity in ID8-OR, ID8-PB, and ID8-P cells, normalized to ID8-OR. **(i)** HR activity in HGS2-OR and HGS2 cells, normalized to HGS2-OR. **(j)** HR activity in ID8-Scr, ID8-Rad52KO#1, and ID8-Rad52KO#2, normalized to ID8-Scr. **(k)** SSA activity in HGS2-Scr, HGS2-Rad52KD#1, normalized to HGS2-Scr. (**l**) Representative immunofluorescence images and (**m**) quantitation of γH2AX foci. Scale bar, 10 μm. Data show mean ± standard deviation of at least three independent experiments, with individual dots for each replicate. P-values were determined by two-sided Student’s t-test for two-group comparisons and one-way ANOVA with Šídák’s multiple comparisons test for comparisons among three or more groups.

To assess the impact of genetic depletion of *Rad52* on DNA damage accumulation, we treated ID8-Scr, ID8-KO#1, and ID8-KO#2 cells with 5 μM olaparib for 72 hours and then counted γH2AX foci. We defined γH2AX positivity as ≥5 foci per nucleus and found that the percentage of γH2AX-positive cells was significantly higher in *Rad52* KO cells than in Scr cells (Fig. 4e, 4f). This finding was corroborated by Western blot data, which demonstrated that increased γH2AX expression in *Rad52* knockout cells was exclusively observed after olaparib treatment (Supplementary Fig. 6). Collectively, these findings suggest that genetic depletion of *Rad52* reduces both SSA and HR activity and is associated with increased accumulation of DNA damage after olaparib treatment.

### *Polq* expression is upregulated in *Rad52* knockout cells treated with olaparib

To investigate the effect of *Rad52* depletion on expression of DNA repair-related genes, we performed bulk RNA sequencing (RNA-seq) of the ID8-KO#1 and Scr cells after treatment with 1 μM olaparib or vehicle for 24 hours (Fig. 5a). Principal component analysis revealed that the transcriptomic profiles were clearly separated among the four experimental groups (Fig. 5b). Differentially expressed genes were defined as those with a log_2_ fold change ≥ 0.5 and adjusted p-value < 0.05. When comparing *Rad52* KO#1 to Scr control after olaparib treatment, 914 genes were upregulated, and 1038 genes were downregulated (Fig. 5c). Among genes involved in DNA repair, *Polq*, *Xrcc3*, and *Fen1* were upregulated, whereas *Rad51c, Pold4,* and *Brcc3* were downregulated in *Rad52* KO#1 cells treated with olaparib. In the heatmap, genes were ranked based on the average Z-score of mRNA abundance in Scr cells treated with olaparib (Fig. 5d). Among these, *Polq* expression was consistently higher in KO#1 cells than in Scr cells. Furthermore, in a comparison across the four groups based on Z-scores, olaparib-induced upregulation of *Polq* was observed only in KO#1 cells (Fig. 5e). Finally, qRT-PCR analysis confirmed that *Polq* expression was ∼1.61-fold higher in olaparib-treated KO#1 cells than in olaparib-treated Scr cells (Fig. 5f). Taken together, these findings suggest that *Polq* is upregulated in *Rad52* knockout cells after olaparib treatment as a compensatory response to the loss of alternative DNA repair pathways.

**Figure 5.**
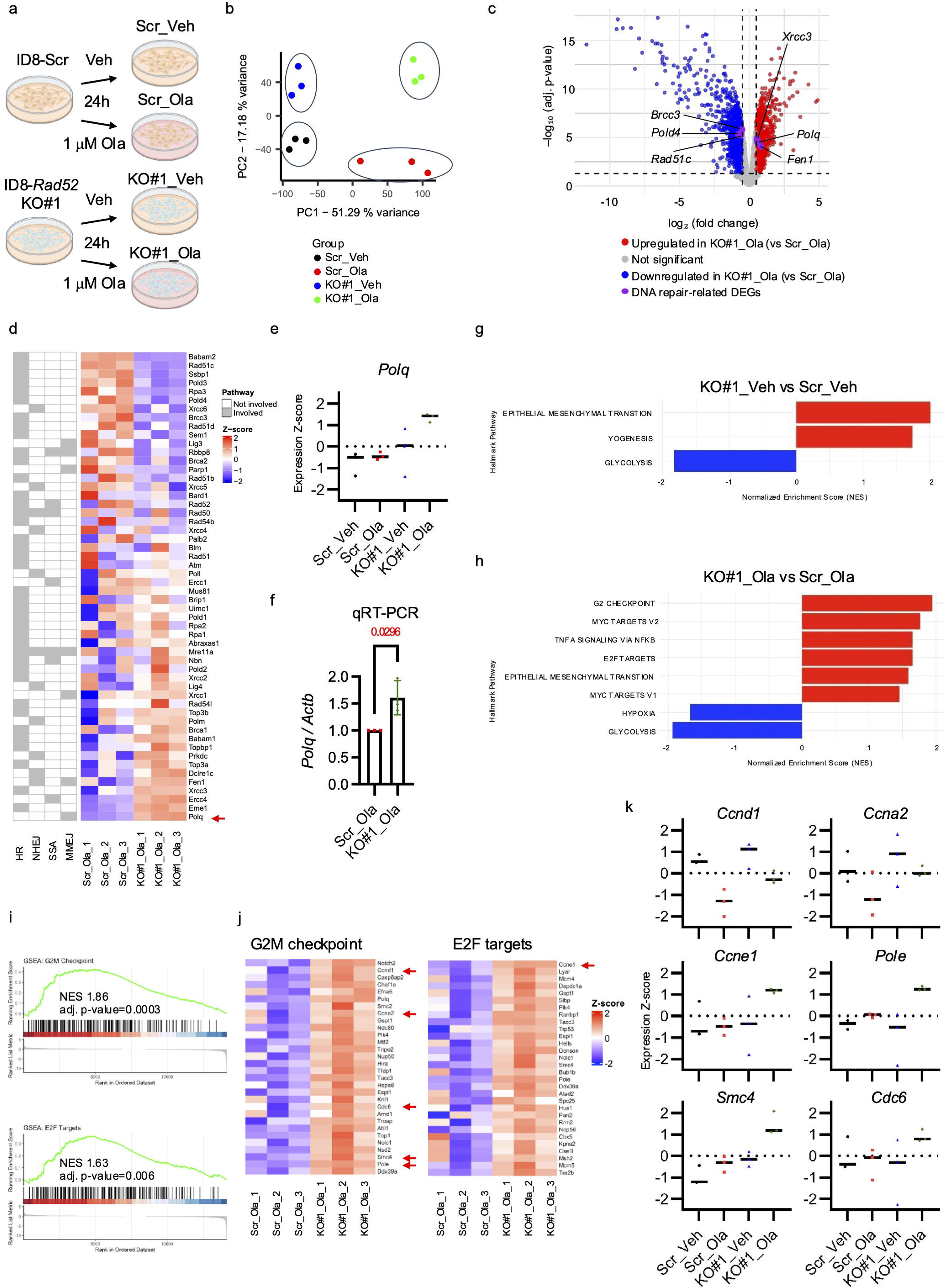
RNA-seq reveals *Polq* upregulation and aberrant activation of the G2M checkpoint and E2F target pathways in *Rad52*-deficient ID8-OR cells treated with olaparib. **(a)** Schematic illustration of the cell lines and treatment groups. **(b)** Principal component analysis of the four experimental groups. **(c)** Volcano plot shows differentially expressed genes between ID8-KO#1 and Scr cells treated with olaparib. Differentially expressed DNA repair-related genes are labeled. **(d)** Heatmap showing Z-score expression values of DNA repair–related genes, ranked in descending order based on expression in Scr cells treated with olaparib. Genes involved in specific pathways are labeled. **(e)** Z-score of *Polq* expression in Scr and ID8-KO#1 cells treated with olaparib or vehicle. **(f)** qRT-PCR analysis of *Polq* expression, normalized to b-actin and presented relative to the expression in Scr cells treated with olaparib. **(g, h)** Gene set enrichment analysis of Hallmark pathways between ID8-KO#1 and Scr control treated with **(g)** vehicle or **(h)** 1 μM olaparib for 24 hours. Enrichment bar plot shows normalized enrichment scores. Pathways with adjusted p-value <0.05 were considered significant. **(i)** Enrichment plots of G2M checkpoint and E2F targets pathways. **(j)** Heatmap shows top 30 leading-edge genes of G2M checkpoint and E2F Targets pathways with the highest log fold-change values in Rad52-KO#1 versus Scr control cells treated with olaparib. **(k)** The dot plot shows the Z-scores of *Ccnd1*, *Ccna2*, *Ccne1*, *Pole*, *Smc4*, *and Cdc6* (red arrows in **j**) in the four conditions (ID8-Scr and ID8-*Rad52*KO#1 cells treated with vehicle or olaparib). Data are based on RNA-seq analysis. Bars indicate the mean of three independent experiments, with individual dots representing each replicate.

### RAD52 knockout leads to alterations in genes involved in cell cycle regulation

We next performed Gene Set Enrichment Analysis with the Hallmark gene sets from the Molecular Signatures Database to explore broader biological changes beyond DNA repair. In the vehicle-treated condition, *Rad52* KO#1 cells exhibited positive enrichment of the epithelial mesenchymal transition and myogenesis pathways and negative enrichment of glycolysis pathways (Fig. 5g). Upon olaparib treatment, additional pathways were positively enriched, including G2M checkpoint, E2F targets, MYC targets (V1 and V2), and TNFα signaling via NFκB, whereas hypoxia and glycolysis pathways were negatively enriched (Fig. 5h). Given the well-established link between DNA repair and cell cycle regulation, we focused on enrichment of the G2M checkpoint and E2F target pathways in the olaparib-treated cells. These pathways were more enriched in olaparib-treated *Rad52* knockout cells than in olaparib-treated Scr control cells (Fig. 5i). Next, we analyzed the top 30 leading-edge genes with the highest log fold change values within these two pathways and confirmed that these genes were upregulated (Fig. 5j). We then compared their expression in Scr and *Rad52* KO#1 cells in vehicle- and olaparib-treated conditions and selected genes that exhibited characteristic expression changes (Fig. 5k, Supplementary Fig. 7). *Ccnd1* (cyclin D1), essential for the G1/S transition, and *Ccna2* (cyclin A2), required for both the G1/S and G2/M transitions^54,55^, were lower after olaparib treatment than after vehicle treatment in both *Rad52* KO#1 cells and Scr controls. Interestingly, the extent of reduction induced by olaparib tended to be smaller in KO cells than in Scr cells. In contrast, the following genes were upregulated upon olaparib treatment in *Rad52* KO cells but not in Scr controls: *ccne1* (cyclin E1), an E2F target crucial for G1/S progressio^56^, *pole* (Polymerase Epsilon), a catalytic subunit of DNA polymerase ε responsible for leading-strand synthesis^57^, *smc4* (SMC4), a condensin complex component essential for chromosome condensation^58^, and *cdc6* (CDC6), a licensing factor for DNA replication origins^59^.

### Small molecule inhibition of RAD52 restores PARP inhibitor sensitivity in *Brca2*-deficient PARP inhibitor-resistant ovarian cancer cell lines

To investigate whether pharmacological inhibition of RAD52 could overcome PARP inhibitor resistance, we tested the selective small-molecule RAD52 inhibitor D-I03, which blocks RAD52-dependent SSA and D-loop formation^60^. D-I03 significantly decreased the olaparib IC₅₀ in ID8-OR and HGS2-OR cells (Fig. 6a). To determine whether olaparib synergized with D-I03, we used the BLISS model, which assesses the expected combined effect assuming independent drug action, and the HSA model, which compares the observed combination effect to the more potent single-agent effect^61^. In both models, scores greater than 0 indicate synergy and scores less than 0 suggest antagonistic effects. Synergistic interactions between olaparib and D-I03 were observed in both cell lines (Fig. 6b). Similar results were obtained with Mitoxantrone, which blocks the RPA-RAD52 interaction and has been used as RAD52 inhibitor *in vitro* (Supplementary Fig 8)^62^. These findings suggest that RAD52 inhibitors synergize with olaparib.

**Figure 6.**
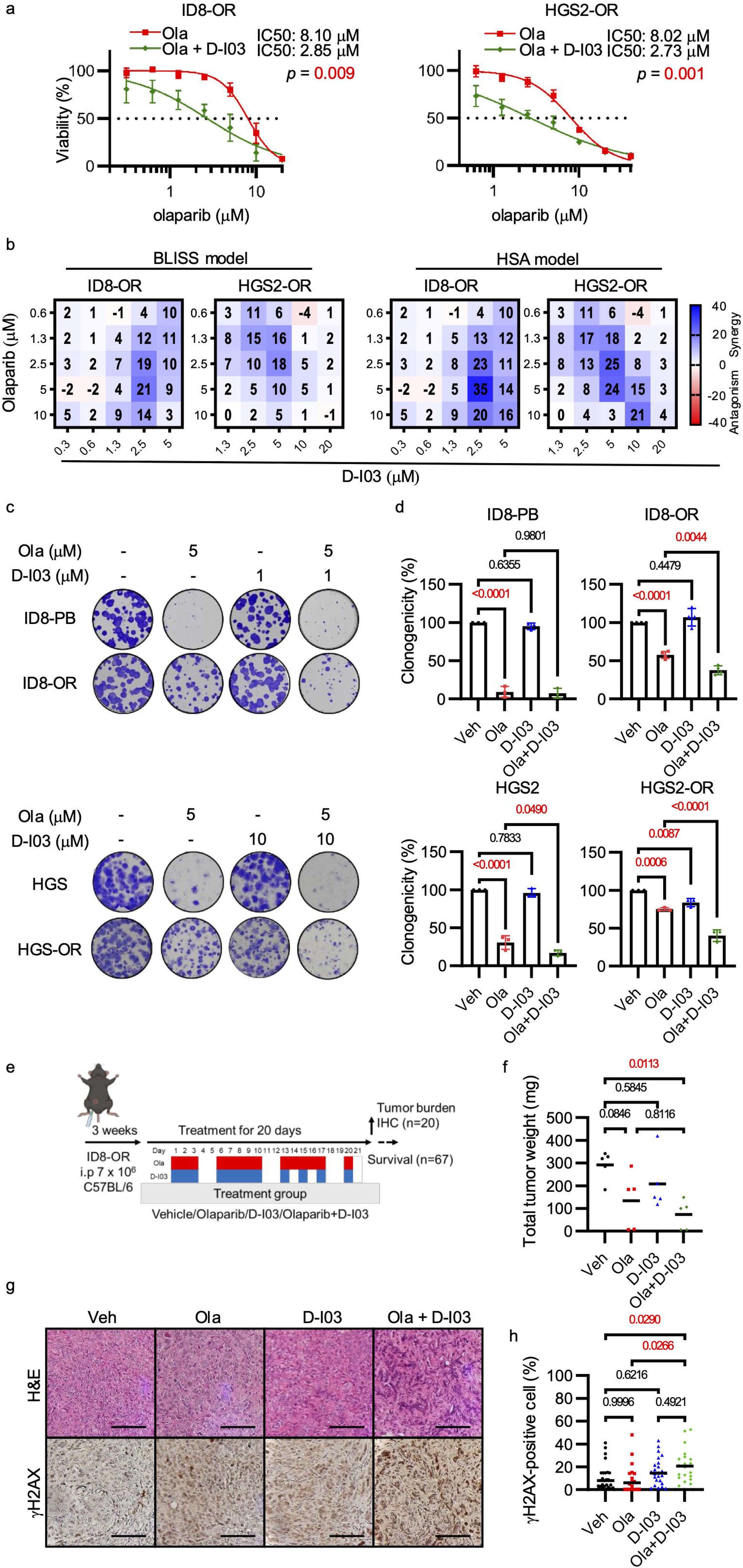
Selective RAD52 inhibitor restores PARP inhibitor sensitivity in *Brca2*-deficient, PARP inhibitor–resistant ovarian cancer cells *in vitro* and *in vivo*. **(a)** Viability assay of cells treated with olaparib alone or in combination with D-I03 for 5 days. **(b)** BLISS and HSA synergy map based on the data in (**a**). **(c)** Clonogenicity assay of ID8-PB, ID8-OR, HGS2, and HGS2-OR cells treated with the indicated concentrations of olaparib and D-I03. **(d)** Quantification of clonogenic assay results. **(e)** Schematic illustrating the ID8-OR xenograft mouse model. **(f)** Tumor weights at the end of treatment. **(g)** Representative images of H&E, immunohistochemistry of γH2AX. Scale bar, 100 μm. **(h)** Quantification of γH2AX-positive nuclei in intraperitoneal tumor tissues. Data show mean ± standard deviation of at least three independent experiments, with individual dots for each replicate. P-values were determined by two-sided Student’s t-tests for two-group comparisons and one-way ANOVA followed by Šídák’s multiple comparisons test for comparisons among three groups.

Next, we assessed clonogenicity after treatment. ID8-OR and HGS2-OR cells formed fewer clones after treatment with 5 μM olaparib than after vehicle treatment (Fig. 6c, 6d). The reduction in clonogenicity was even greater in cells treated with olaparib plus D-I03. In the parental ID8-PB and HGS2 cells, olaparib alone also significantly decreased clonogenicity.

To evaluate the *in vivo* efficacy of D-I03, ID8-OR cells (7 × 10⁶) were intraperitoneally inoculated into C57BL/6 mice. Beginning three weeks after tumor cell inoculation, olaparib (75 mg/kg, oral gavage), D-I03 (25 mg/kg, intraperitoneal), the combination of both drugs, or vehicle were administered for 20 days (Fig. 6e). Two mice from the D-I03 and combination groups died shortly after treatment initiation without detectable tumors, suggesting drug-related toxicity (Supplementary Fig. 8). At the end of the treatment period, mice that received D-I03 plus olaparib had smaller tumor weight (Fig. 6f) than mice treated with vehicle. Tumors from mice treated with D-I03 plus olaparib had more γH2AX-positive cells than tumors from mice in the vehicle and olaparib monotherapy groups (Fig. 6g, 6h). The combination treatment prolonged survival relative to the vehicle group, extending the 25% survival time by 7 days (Supplementary Fig.8). Collectively, these findings demonstrate that pharmacological inhibition of RAD52 restores PARP inhibitor sensitivity in *Brca2*-deficient, PARPi-resistant ovarian cancer models, both *in vitro* and *in vivo*, by promoting accumulation of DNA damage.

## Discussion

Here, we provide evidence that RAD52 is a promising therapeutic target to overcome PARPi resistance in *Brca2*-deficient ovarian cancer. Elevated RAD52 expression was associated with poor prognosis in HGSC. Additionally, both genetic and pharmacologic inhibition of RAD52 re-sensitized *Brca2*-deficient, PARPi-resistant cells *in vitro* and *in vivo*. Mechanistically, RAD52 depletion impaired both SSA and HR, leading to accumulation of DNA double-strand breaks after PARPi treatment. Furthermore, RAD52 upregulation observed in PARPi-resistant models suggests an adaptive mechanism that partially contributes to resistance, as increased SSA and HR activity were associated with elevated RAD52 expression. Transcriptomic analysis further revealed that PARPi treatment induced *Polq* expression in *Brca2*- and *Rad52*-deficient cells, suggesting a compensatory reliance on microhomology-mediated end joining (MMEJ) as a survival pathway. These findings establish RAD52 as a mechanistically and therapeutically relevant target for overcoming acquired PARPi resistance in *BRCA2*-deficient ovarian cancer.

Our findings provide further insight into the role of RAD52 in supporting homologous recombination (HR) in *BRCA2*-deficient ovarian cancer cells and suggest that it contributes to PARPi resistance. In our study, knockdown of *Rad52* reduced HR activity in HGS2-OR cells, consistent with studies showing that *RAD52* deletion in *BRCA2*-deficient breast cancer models further impairs HR^34^. Moreover, HGS2-OR cells exhibited higher HR activity than their parental HGS2 cells, consistent with reports linking RAD52 to platinum resistance in ovarian cancer^63^. This finding suggests that RAD52-mediated HR is restored in the absence of *Brca2* reversion mutations. Although the C-terminal domain of mammalian RAD52 can bind RAD51, it cannot load RAD51 onto DNA as efficiently as BRCA2^64^. Instead, RAD51–RAD52 mixed filaments can mediate HR, albeit inefficiently^64^. Mechanisms reported to promote RAD52 activity include post-translational modifications of RAD52 (e.g., SUMOylation^65^, phosphorylation^66^, and acetylation^67,68^), interactions with phosphorylated RPA^69^, and cofactors such as DSS1^70^. Given the modest extent of HR upregulation in HGS2-OR cells, these factors may collectively enhance the contribution of RAD52 to HR. In ID8-OR cells, deletion of *Rad52* also reduced HR activity. However, the HR activity in ID8-OR cells remained comparable to that of the parental ID8-PB cells and was lower than that in the HR-proficient ID8-P model. These findings suggest that, in ID8-OR cells, RAD52 primarily supports residual HR activity rather than substantially restores HR.

We also elucidated the contribution of RAD52-mediated single-strand annealing (SSA) to PARPi resistance. In mammalian cells, SSA is a well-established function of RAD52^35^. Similar to our observations with HR, knockdown of *Rad52* reduced SSA activity in HGS2-OR cells. Moreover, SSA activity was higher in HGS2-OR cells than in their parental HGS2 cells, suggesting that increased SSA, along with HR, contributes to PARPi resistance in this model. In ID8-OR cells, *Rad52* knockout similarly reduced SSA activity. Although SSA activity did not differ between ID8-OR and their parental ID8-PB cells, it was higher than that in the HR-proficient ID8-P cells, indicating a greater reliance on SSA in *Brca2*-deficient cells^71^.

Taken together, our findings reveal two distinct modes of RAD52 involvement in *BRCA2*-deficient, PARPi-resistant cells. In HGS2-OR cells, RAD52 promotes resistance through both SSA and HR, whereas in ID8-OR cells, RAD52 primarily supports cell survival via SSA and residual HR activity. Notably, RAD52 inhibition sensitized both models to PARPi, highlighting RAD52-dependent repair as a key therapeutic vulnerability. Given the pronounced intertumoral heterogeneity and diversity of resistance mechanisms in ovarian cancer, precise patient stratification is essential^72,73^. Factors such as BRCA mutation status, RAD52 expression levels, and functional biomarkers including SSA and HR activity should be incorporated to identify patients most likely to benefit from RAD52-targeted therapy.

Our RNA-seq analysis provided several insights into the effects of *Rad52* knockout on PARPi-resistant, *Brca2*-deficient cells. *Rad52* loss triggered *Polq* upregulation after PARPi treatment, suggesting a compensatory shift toward the MMEJ pathway. POLQ is under active investigation as a therapeutic target in *BRCA*-deficient tumors^46,74-76^. Both SSA and MMEJ require 5′ to 3′ end resection to generate 3′ overhangs, followed by annealing of homologous sequences, which results in deletions and renders these pathways inherently mutagenic. In HeLa cells, RAD52 suppressed POLQ-mediated MMEJ activity, thereby preventing its premature activation during the G2/M phase^77^. Although the mechanisms determining pathway choice between SSA and MMEJ are poorly understood, our results indicate that targeting RAD52 forces tumor cells to adopt a survival strategy mediated by POLQ, thereby increasing their dependence on POLQ. Thus, in addition to highlighting RAD52 as a therapeutic target, our findings suggest that combined inhibition of RAD52 and POLQ is a promising approach for treatment optimization, particularly in PARPi-resistant *BRCA*-deficient tumors^46^.

In *Rad52*-wild-type cells, olaparib treatment led to a marked reduction in *Ccnd1* and *Ccna2*, consistent with a protective growth-suppressive response^78^. In contrast, Rad52-deficient cells showed only a modest reduction in these genes and instead showed upregulation of *Ccne1*, *Pole*, *Cdc6*, and *Smc4*, indicating aberrant activation of replication and mitotic programs despite unresolved DNA damage^57-59^. This shift from a growth-suppressive to a proliferative program likely exacerbates replication stress and genomic instability in the presence of a PARP inhibitor, providing a mechanistic basis for the heightened sensitivity observed in *Rad52*-deficient cells.

A key limitation of our paper is that the RAD52 inhibitor D-I03 demonstrated acute toxicity in mice. This likely limited the achievable dose and shortened the treatment duration, contributing to the modest survival benefit observed. These findings underscore the need for safer and more selective next-generation inhibitors.

In conclusion, our study provides compelling preclinical evidence that RAD52 is a viable therapeutic target in *BRCA2*-deficient, PARPi-resistant ovarian cancer. By disrupting residual repair pathways and exposing compensatory vulnerabilities such as POLQ dependency, RAD52 inhibition offers a rational strategy to overcome acquired resistance. The development of potent, selective, and safe RAD52 inhibitors, ideally combined with agents targeting parallel repair pathways, is a critical next step toward clinical application.

## Methods

### Ethics statement

All mice were maintained in compliance with the guidelines approved by the Institutional Animal Care and Use Committee at Washington University in St. Louis (23-0256). A 12-hour light/12-hour dark cycle was maintained, with temperatures between 65 °F and 75 °F and humidity at 40–60%. Every effort was made to ensure the animals did not experience unnecessary discomfort or distress. Mice were euthanized by cervical dislocation.

### Tissue microarray

A previously constructed tissue microarray comprising high-grade serous ovarian carcinoma (HGSC) samples from patients treated at the University of Colorado was provided by the GFTB (COMIRB #17-7788)^79,80^. Slides were immunohistochemically stained for RAD52. Cases with tumor microarray cores containing less than 5% tumor cells, as well as cases derived from recurrent tumors or those treated with neoadjuvant chemotherapy, were excluded from the analysis (Supplementary Fig. 1, Supplementary Table 1). Tumor regions were selected under the supervision of a pathologist, and RAD52 expression was quantified by using the H-score method (range: 0–300)^81^ with QuPath software (Ver. 0.5.1). Patients were stratified into "Low" (H-score <168) and "High" (H-score ≥168) expression groups based on the median value. Survival differences were evaluated by using the log-rank test, with analysis focused on the four-year time point from diagnosis.

### Clinical prognosis analysis from public database

The Cancer Genome Atlas Ovarian Serous Cystadenocarcinoma dataset (PanCancer Atlas) was accessed through cBioPortal (http://www.cbioportal.org) to obtain mRNA data from ovarian cancer patients. This dataset was last accessed on 17 July 2025. For analysis, only *TP53*-mutated samples were selected. mRNA abundance was quantified with RSEM (Batch normalized from Illumina HiSeq_RNAqV2). mRNA abundances were then divided into high and low groups based on the median. Kaplan-Meier analysis was performed to assess the overall survival of patients with high and low *RAD52* mRNA expression. Survival differences were evaluated by using the log-rank test, with analysis focused on the four-year time point from diagnosis.

### Cell lines

ID8-OR and HGS2-OR cell lines were established by co-author Benjamin G. Bitler and colleagues from ID8-PB (*Trp53*^−/−^, *Brca2*^−/−^) and HGS2 (*Trp53*^−/−^, *Brca2*^−/−^, *Pten*^−/−^), respectively, cells as previously described^82,83^. ID8-PB and ID-PB-OR cells were cultured in DMEM (Sigma Aldrich, MO, #D6429) supplemented with 4% fetal bovine serum (FBS), penicillin/streptomycin (Gibco, #15140122), and 1× ITS (Sigma Aldrich, MO)^84^. HGS2 and HGS2-OR were cultured in DMEM/F12 (Gibco, #11320033) supplemented with 10% FBS, penicillin/streptomycin, and 1× ITS (Gibco, #41400-045)^49^. All cells were cultured at 37 °C in 5% CO_2_.

### Next-generation sequencing-based genotyping

The target region was PCR amplified (PCR1) with primers (gene-specific primer sequences below) appended with 5′-CACTCTTTCCCTACACGACGCTCTTCCGATCT-3′ on the forward primer and 5′-GTGACTGGAGTTCAGACGTGTGCTCTTCCGATCT-3′ on the reverse primer to allow unique indexes and Illumina P5/P7 adapter sequences to be added in a second round PCR. PCR amplifications were performed with SuperFi II 2x Platinum Green master mix (ThermoFisher) according to the manufacturer’s protocol. Indexing of the PCR1 product was performed by using 0.1X volume from PCR1 with indexing primers (0.1 μM final concentration for each) and melting at 98 °C for 2 min, followed by five cycles of 98 °C for 15 s, 60°C for 15 s, and 72 °C for 40 s. We generated 2 × 250 reads with the Illumina MiSeq platform at the Center for Genome Sciences and Systems Biology (Washington University). The extracted FASTQ files were analyzed with a Python-based alignment script.

Gene specific primers: mouse Brca2_exon3 forward 5’- CCTTTGGCAACCCTGTTCTT -3’ and reverse 5’- CCACCTGTTTCTGCCTCTCT -3’; mouse Brca2_exon11 forward 5’-GGCTGTCTTAGAACTTAGGCT -3’ and reverse 5’- TGTTGGATACAAGGCATGTAC -3’. (Supplemenrtary file).

### Western blots

Cells were harvested and lysed in RIPA buffer (Cell Signaling, MA, #9806S) with Protease Inhibitor Cocktail (Sigma Aldrich, #P8340) and Phosphatase Inhibitor Cocktail (Sigma Aldrich, #P5726). Cell lysates were separated by SDS-PAGE and immunoblotted with primary antibody and then secondary antibody. Chemiluminescent horseradish peroxidase (HRP) substrate (Millipore, MA, #WBKLS0100) was used for development, and the ChemiDoc Imaging System (Bio-Rad, CA) was used for detection. Densitometric analysis was performed in Image Lab software (BioRad). The following antibodies and dilutions were used: RAD52 (Origene, MD, TA890144, 1:2000), γH2AX (Abcam, Cambridge, UK, ab26350, 1:1000), β-actin (Invitrogen, MA, USA, PA1-183-HRP, 1:2500), Anti-mouse IgG, HRP (Cytiva, MA, USA, NA931V, 1:5000), Anti-rabbit IgG, HRP (Cytiva, NA934V, 1:5000).

### qRT-PCR

Total RNA was extracted with the RNeasy Protect Mini Kit (Qiagen, MD, #74124) and then reverse transcribed to cDNA with the QuantiTect Reverse Transcription Kit (Qiagen, #205313) according to the manufacturer’s instructions. cDNAs were analyzed by q-PCR using Fast SYBR Green Master Mix (Applied Biosystems, CO, USA, #4385612) on a 7500 Fast Real-Time PCR system (Applied Biosystems), and data were normalized to b-actin. Relative gene expression was analyzed by the 2 ^-ΔΔCt^ method. The primers used were as follows: mouse *Rad52* forward 5’-GAAGGTGTGTTATATTGAAGGTCATCG-3’ and reverse 5’- AATCCACATTTTGCTGGGTGATGG-3’, mouse *Polq* forward 5’-GTCACAGTCACACACGGCTA-3’ and reverse 5’-CATTGGGAACCGGCCTTTTG-3’, mouse *Actb* forward 5’- TTACTGCTCTGGCTCCTAGC-3’ and reverse 5’-CAGCTCAGTAACAGTCCGC-3’ (Supplemenrtary file).

### CRISPR/CAS9 knockout

Predesigned guide RNA for mouse *Rad52* and the CRISPR-Cas9 system were used according to the manufacturer’s protocol (Integrated DNA Technologies, IA). The crRNA targeting sequence GTGTTATATTGAAGGTCATC (Mm.Cas9.RAD52.1.AE) and tracrRNA labeled with ATTO 550 dye were annealed to form guide RNA. Then, guide RNA and CAS9 protein were combined to form ribonucleoprotein complexes. Sub-confluent ID8-OR cells were transfected with Lipofectamine RNAiMAX (Invitrogen, #13778150). The top 10% of cells with the highest ATTO fluorescence intensity were single-cell sorted by flow cytometry. Single cell clones were isolated, screened by Guide-it Genotype Confirmation kit (Takara Bio, Kusatsu, Japan, #632611), and validated by Western blot, qRT-PCR, and next-generation amplicon sequencing (GENEWIZ, NJ, USA, Amplicon-EZ,) with the following primers: mouse *Rad52* forward 5’- GCTCTGTACACCCAGCTACC-3’ and reverse 5’-AGTGTGCCCAGCCATTGTAA-3’. The alleles identified by next-generation sequencing at frequencies >5% are shown (Supplementary file).

### RNA interference, lentivirus, and plasmid transfection

*Rad52*-silenced cell lines were generated with a pool of three short hairpin RNA (shRNA) lentiviral particles targeting mouse *Rad52* (Santa Cruz, TX, USA, sc-37400-V), according to the manufacturer’s instructions. HGS2-OR cells were transfected with lentiviruses and 5 μg/mL polybrene (Santa Cruz, sc-134220) and selected with 8 μg/mL puromycin (Santa Cruz, sc-108071) for three days. Cells transfected with scrambled shRNA (Santa Cruz, sc-108080) were used as controls.

### Immunofluorescence

Immunofluorescence was performed as previously described^52^. Briefly, cells were seeded on sterilized cover slips in a 12-well plate. After treatment with olaparib for 72 hours, cells were washed with cold PBS++ (Leinco Technologies, MO, # P364), fixed with 2% paraformaldehyde (Thermo Fisher, MA, #J19943-K2) in PBS++ for 10 minutes, permeabilized with 0.2% Triton X-100 in PBS (Sigma-Aldrich, #T8787) for 20 minutes, and then washed and blocked (30 minutes) with staining buffer (PBS, 0.5% BSA, 0.15% glycine, and 0.1% Triton-X-100). Cells were incubated overnight at 4 °C with primary antibodies in staining buffer and then stained with secondary antibodies at room temperature for 2 hours. Slides were mounted in ProLong Gold (Invitrogen, MA, #P36930). The images were obtained on a confocal microscope (DMI4000B, Leica, Wetzlar, Germany) equipped with a 63x oil immersion objective lens. Images were acquired in 512 × 512 pixel format using Leica Application Suite X and analyzed with ImageJ software (version 1.52a)^85^. Color images were converted to 8-bit grayscale, and thresholds were set appropriately and consistently. γH2AX foci were counted by using the ‘analyze particles’ function. At least 100 cells were analyzed for each treatment group. Cells with 5 or more nuclear γH2AX foci were defined as positive, and the proportions of positive cells were compared. The following antibodies and dilutions were used: γH2AX (Millipore Sigma, 05-636, 1:500), Anti-mosue IgG, Alexa Fluor 555 conjugate (Cell Signaling Technology, 4409, 1:500) (Supplementary file).

### Viability assay

Cells were seeded in 96-well plates overnight and then treated with the indicated drugs for five days. Media with drug was replaced on the third day of treatment. Cells were stained and fixed with crystal violet solution (0.05% Crystal Violet [Fisher, Cat. #C581-25], 1% Formaldehyde [Sigma-Aldrich, Cat. #252549], 1X PBS [Corning, Cat. # 21040-CM], 1% Methanol [Sigma-Aldrich, Cat. #1003657780] in dH2O), then washed in 70% ethanol (Fisher Scientific, Cat. #04355223). Viability was evaluated by measuring absorbance at 570 nm. Detailed information for each cell line is provided in Supplementary file.

### Clonogenicity assay

Cells were seeded in 12-well plates and incubated overnight. Then, cells were treated with the indicated drugs, and drug-containing media was refreshed every 3 days for a total of 7–8 days. Colony formation was assessed as described for viability assays. Detailed conditions for each cell line are provided in Supplementary file.

### Drug synergy analysis

Cells were seeded in 96-well plates overnight and then co-treated for 5 days with olaparib and D-I03. Cell viability was quantified by crystal violet absorbance as described above. Bliss and HSA scores were analyzed by Combenefit ver. 2.021 (https://sourceforge.net/projects/combenefit/)^86^ and visualized in Graphpad Prism 9.

### DNA repair assay

SSA activity was evaluated with the hprtSAGFP-based reporter system, and HR activity was evaluated with the pDR-GFP-based reporter system, as previously reported^87^. Briefly, the reporter constructs were hprtSAGFP (Addgene, MA. USA, plasmid #41594; http://n2t.net/addgene:41594; RRID:Addgene_41594), pDR-GFP (Addgene plasmid #26475; http://n2t.net/addgene:26475; RRID:Addgene_26475), and pCBASceI (Addgene plasmid #26477; http://n2t.net/addgene:26477; RRID:Addgene_26477). These plasmids were gifts from Dr. Maria Jasin^88,89^. Cells were seeded in 12-well plates and incubated overnight. Then, cells were co-transfected with either hprtSAGFP or pDR-GFP together with pCBASceI by using Lipofectamine 3000 (Invitrogen, #L3000008). After 24 –48 hours, cells were harvested, and the percentages of GFP-positive cells were determined by flow cytometry (CYTEK, CA, USA, AURORA) using B1 channel and analyzed (CYTEK, Spectro Flo, Version 3.3.0). The percentages of GFP-positive cells transfected with only hprtSAGFP or pDR-GFP (without pCBASceI) were subtracted as background^87^.

### Mouse experiments

For experiments with ID8-Scr or ID8-KO#1 cells, 5 × 10⁶ cells were intraperitoneally injected into 6- to 8-week-old female C57BL/6 mice (Jackson Laboratory). One week after implantation, mice received daily treatment with olaparib (50 mg/kg, oral gavage) or vehicle for four weeks^84^. Eight mice per group were euthanized the day after the final treatment, and tumor burden was evaluated by measuring total tumor weight (excluding omentum), omental weight, the number of peritoneal tumor nodules larger than 1 mm in diameter, and ascites volume.

For experiments involving RAD52 inhibitors, 7 × 10⁶ ID8-OR cells were intraperitoneally injected into C57BL/6 mice. Three weeks after implantation, mice were treated for 20 days with olaparib (75 mg/kg, oral gavage)^90^, D-I03 (25 mg/kg, intraperitoneal injection), olaparib plus D-I03, or vehicle. The day after treatment completion, five mice per group were euthanized, and tumor burden was assessed by measuring total tumor weight.

In both experimental settings, tumors were subjected to immunohistochemical analysis for γH2AX and RAD52. Survival cohorts were monitored after treatment completion in accordance with the Guide for the Care and Use of Laboratory Animals (8th edition, National Research Council, 2011), and all procedures were approved by the Institutional Animal Care and Use Committee of Washington University in St. Louis. All animals were monitored daily for clinical signs of distress and changes in body weight. Mice in which no macroscopic tumor engraftment was observed at necropsy were excluded from survival analysis as non-engrafted cases.

### RNA-seq

Total RNA was extracted from ID8-Scr control cells and *Rad52* KO#1 treated with 1 μM olaparib or vehicle for 24 hours. RNA integrity was assessed by using an Agilent Bioanalyzer or 4200 Tapestation, and samples with RNA integrity number > 8.0 were used for library preparation. Polyadenylated RNA was isolated with oligo-dT beads (mRNA Direct kit, Life Technologies), fragmented at 94 °C, and reverse transcribed into cDNA with SuperScript III reverse transcriptase and random hexamers. After second-strand synthesis, cDNA was end-repaired, A-tailed, ligated to Illumina adapters, and PCR-amplified with dual-index primers. Libraries were sequenced on an Illumina NovaSeq X Plus platform with 2 × 150 bp paired-end reads. Base calling and demultiplexing were performed with Illumina’s bcl2fastq software. Reads were aligned to the Ensembl GRCm39.113 reference genome in STAR (v2.7.11b), and gene-level counts were obtained by using Subread:featureCounts (v2.0.8). Isoform expression was quantified with Salmon (v1.10.0). Sequencing quality metrics, including ribosomal fraction, junction saturation, and read distribution across gene models, were assessed with RSeQC (v5.04). Gene expression normalization was performed in the limma package with the voomWithQualityWeights transformation to stabilize the mean–variance relationship. Principal component analysis was conducted by using the prcomp function from base R (stats package), with centering and scaling. Principal component analysis plots were generated in the ggplot2 package based on voom-transformed expression values. Differentially expressed genes were defined as those with an absolute log₂ fold-change (|log₂FC|) > 0.5 and adjusted *p*-value < 0.05 according to the Benjamini–Hochberg method. A heatmap was generated with voom-transformed Z-scores for DNA repair-related genes in ID8-Scr and ID8-KO#1 cells treated with olaparib. Gene sets for HR and NHEJ were defined based on KEGG pathway entries (hsa03440 and hsa03450, respectively), and SSA and MMEJ gene sets were curated from published studies^91,92^. A full list of genes is provided in Supplementary file. Genes were ordered based on descending average Z-score in the Scr group. Additionally, voom-based Z-score expression values for *Polq* and *Rad51C* were shown across all four experimental groups (Scr and ID8-KO#1 with or without olaparib treatment) to illustrate treatment- and genotype-specific regulation. Gene Set Enrichment Analysis was performed with the Hallmark gene sets from the Molecular Signatures Database. Comparisons between *Rad52* knockout and Scr control cells were conducted in both vehicle- and olaparib-treated conditions. Pathways significantly enriched at an adjusted p-value cutoff of 0.05 were ranked according to their normalized enrichment scores. Enrichment plots were generated for the G2M checkpoint and E2F target pathways. For these pathways, the top 30 leading-edge genes with the highest log fold change values were selected, and their voom-transformed expression values in the olaparib-treated cells were visualized as a heatmap.

### Statistics

Two-sided Student’s *t*-test was used for comparisons between two groups. For comparisons among three or more groups, one-way ANOVA followed by Šídák’s test was performed. Selected pairwise comparisons were adjusted for multiple testing by using the Šídák method. All data are presented as mean ± standard deviation unless otherwise stated. GraphPad Prism 9 was used for all analyses.

## Data availability

RNA-seq data for ID8-OR-Rad52 and Scr control with vehicle and olaparib are deposited into GEO under accession number GEO: GSE302064. All code used for RNA-seq data processing and analysis is available upon reasonable request.

## Author contributions

YO conceptualized experiments, conducted experiments, acquired and analyzed data, and drafted the manuscript. VG, BF, MA, and PB acquired data and contributed to manuscript editing. PT performed RNA-seq data processing and analysis. LS analyzed and confirmed pathology of samples. BS, LK, CM, AH, IH, PHT, DG, and MP provided feedback and contributed to manuscript editing. KH performed TMA imaging and data acquisition. PV, JK, BB, and MM advised on experimental design and contributed to manuscript editing. BB also provided cell lines (ID8-OR and HGS2-OR) and TMA samples. DK provided funding and reagents, conceptualized experiments, analyzed data, and revised the manuscript.

## Acknowledgements

We especially acknowledge the patients who generously donated their cancer specimens, making this work possible. We thank Deborah Frank, PhD, for manuscript editing. We acknowledge support from the Gushinkai Fellowship (Support for Young Study Abroad). Imaging of the TMA was performed at the Hope Center Alafi Neuroimaging Lab at Washington University School of Medicine, supported by the NIH Special Instrumentation Grant (S10 OD032121). We also thank the Alvin J. Siteman Cancer Center at Washington University School of Medicine and Barnes-Jewish Hospital, and the Institute of Clinical and Translational Sciences (ICTS) at Washington University in St. Louis, for the use of the Flow Cytometry Core (cell sorting), the Histology Innovation and Service Center (immunohistochemistry), and the Genome Technology Access Center (RNA sequencing), which are supported in part by the NCI Cancer Center Support Grant P30 CA091842. This research was supported by NCI R01CA243511 (DK) and NCI R21CA210210 (DK).

---

**Supplementary Figure 1. Flow diagram of patient selection for the tissue microarray (TMA).** A total of 137 patients with high-grade serous ovarian cancer were initially considered. From this cohort, 26 patients were excluded due to insufficient tumor volume (tumor area <5% of each core), 20 patients whose samples were from recurrent tumors were excluded, and 19 patients were excluded because they had received neoadjuvant therapy, which could potentially alter the molecular profile. This resulted in a final cohort of 72 patients for the analysis.

**Supplementary Figure 2. Evaluation of *Brca2* deletion using next-generation sequencing (a)** DNA sequencing of mouse *Brca2* exon 3 in ID8-P, ID8-PB, and ID8-OR cell lines. **(b)** DNA sequencing of mouse *Brca2* exon 11 in HGS2 and HGS2-OR cell lines.

**Supplementary Figure 3. Generation and validation of *Rad52* knockout ID8-OR cells. (a)** Schematic illustrating the generation of *Rad52*-depleted PARP inhibitor (PARPi) resistant ovarian cancer cells. **(b)** Schematic of the RAD52 domain structure, showing the DNA-, RPA-, and RAD51-binding domains, and a nuclear localization signal (NLS). The CRISPR/Cas9-mediated truncation occurs at amino acid 68. **(c)** Guide RNA target sequence in exon 4 of the mouse *Rad52* gene and the corresponding CRISPR/Cas9 cleavage site. **(d)** Amplicon sequencing of the target region in wild-type (Scramble) and *Rad52* knockout monoclonal cells. **(e)** Quantitative validation of RAD52 knockout by Western blot. Data are presented as mean ± standard deviation from three independent experiments, with individual data points shown for each biological replicate. Statistical significance was assessed by two-sided Student’s t-tests for two-group comparisons and one-way ANOVA followed by Šídák’s multiple comparisons test for comparisons among three groups. KO, knockout; RNP, ribonucleoprotein-based; KD, knockdown; Scr, scrambled; shRNA, short hairpin RNA.

**Supplementary Figure 4. Tumor burden and ascites volume at different time points in mice bearing ID8-Scr or *Rad52*-knockout (ID8-KO#1) tumors from the survival cohort shown in Figure 3. (a)** Representative macroscopic abdominal images at necropsy, performed upon meeting euthanasia criteria in the survival cohort. Scr_Veh and Scr_Ola mice were dissected on day 49, and KO#1_Veh and KO#1_Ola mice were dissected on day 67 after tumor inoculation. The dotted areas indicate disseminated tumor nodules in the peritoneal cavity. Green arrows indicate the omentum. **(b)** Tumor weight (excluding the omentum) and ascites volume measured at the time of euthanasia. Veh, vehicle; Ola, olaparib.

**Supplementary Figure 5. Representative flow cytometry plots from single-strand annealing (SSA) and homologous recombination (HR) reporter assays.**

**(a)** Gating strategy used to exclude dead cells and doublets. **(b–e)** SSA activity assessed by the percentage of GFP-positive cells following co-transfection with hprtSAGFP and pCBASceI plasmids. Background GFP expression was measured following transfection with hprtSAGFP alone. GFP fluorescence was detected in the B1-A channel. Data are shown for **(b)** ID8-OR, ID8-PB, and ID8-P **(c)** HGS2-OR and HGS2 **(d)** ID8-Scr, ID8-*Rad52*KO#1, and ID8-*Rad52*KO#2 and **(e)** HGS2-Scr and HGS2-*Rad52*KD#1. **(f–i)** HR activity assessed using pDR-GFP and pCBASceI plasmids in the same manner. Data are shown for **(f)** ID8-OR, ID8-PB, and ID8-P, and **(g)** HGS2-OR and HGS2 **(h)** ID8-Scr, ID8-KO#1, and ID8-KO#2 **(g)** HGS2-Scr and HGS2-KD#1 **(h)** ID8-OR, ID8-PB, and ID8-P, and **(i)** HGS2-OR and HGS2. **(j)** Negative control confirming the absence of GFP signal in untransfected cells.

**Supplementary Figure 6. Depletion of *Rad52* upregulates γH2AX expression in olaparib-treated cells.** Western blot analysis showing γH2AX expression in ID8-Scramble (Scr) and ID8-*Rad52* knockout (KO#1, KO#2) cells. Proteins were collected from cells treated with 5 μM olaparib or vehicle for 72 hours. γH2AX expression was normalized to β-actin and presented as fold change relative to Scr cells treated with olaparib.

**Supplementary Figure 7. Top 30 leading-edge genes in the G2M checkpoint and E2F pathways in olaparib-treated cells.** Pathways were selected based on the top 30 leading-edge genes with the highest log fold-change values in *Rad52*KO#1 versus Scr control cells treated with olaparib. The dot plots show the Z-scores of these genes across four conditions (ID8-Scr and *Rad52*KO#1 cells treated with vehicle or olaparib). Data are based on RNA-seq analysis. Bars indicate the mean of three independent experiments, with individual dots representing each replicate.

**Supplementary Figure 8. Selective RAD52 inhibitor enhances sensitivity to olaparib in PARP inhibitor–resistant ovarian cancer cells *in vitro* and *in vivo*. (a)** Cell viability matrix showing the combinatorial effects of olaparib with D-I03 or mitoxantrone in ID8-OR cells. **(b)** Cell viability curves for olaparib alone and in combination with 7.5 nM mitoxantrone in ID8-OR cells, based on the data shown in panel **(a)**. Olaparib IC₅₀ values were derived from dose–response curves. **(c)** BLISS and HSA synergy maps based on the data shown in **(a)**. **(d)** Kaplan–Meier survival curves for all mice. An overall log-rank test was followed by pairwise comparisons between selected groups. Median survival (days): Veh, 56.5; Ola, 58; D-I03, 57; Ola + D-I03, 58. The time points corresponding to 25% survival were: Veh, 57 days; Ola, 58 days; D-I03, 60 days; Ola + D-I03, 62 days. **(e)** Kaplan–Meier survival curves excluding suspected drug-related deaths. Median survival (days): Veh, 56.5; Ola, 58; D-I03, 57; Ola + D-I03, 60. The time points corresponding to 25% survival were: Veh, 57 days; Ola, 58 days; D-I03, 60 days; Ola + D-I03, 64 days. P-values determined by Bonferroni-corrected threshold of p < 0.0167 (0.05/3) for multiple comparisons.

