## Supplementary_figure_table for "Targeting RAD52 overcomes PARP inhibitor resistance in preclinical *Brca2*-deficient ovarian cancer model"

**Supplementary Figure 1**

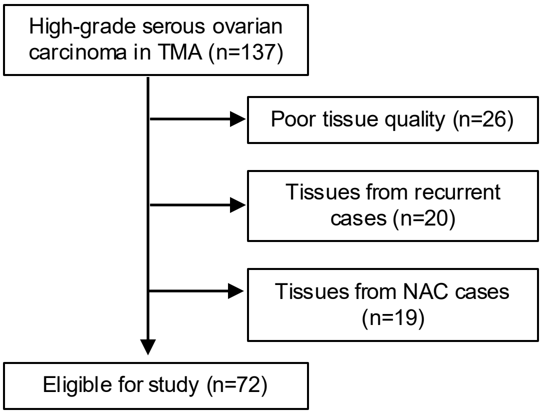

**Supplementary Figure 2**

Site of the *Brca2* mutation

a

ID8-P
*Brca2 wt,*
Refference

NM_001081001.2:c.=, NP_001074470.1:p.(=)

exon2 | exon3

agatgcagcacagcagatttaggaccgataagcctcaattggtttgaggagctttcctca
R C S T A D L G P I S L N W F E E L S S
gaagccccccatacaattctgaacctccggaggaatctgagtataagccccacggttat
E A P P Y N S E P P E E S E Y K P H G Y
gaaccacagctgtttaaaacaccacagaggaatccccctaccatcagtttgcttcaact
E P Q L F K T P Q R N P P Y H Q F A S T
ccaataatgttcaaa
P I M F K

ID8-PB/OR
*Brca2 del*

NM_001081001.2:c.[69T>A;94_219del] NP_001074470.1:p.[Asp23Glu;Phe32_Gln73del]

Allele 1

exon2 | exon3

agatgcagcacagcagaattaggaccgataagcctcaattgg------------------
R C S T A E L G P I S L N W
------------------------------------------------------------
------------------------------------------------tttgcttcaact
F A S T
ccaataatgttcaaa
P I M F K

NM_001081001.2:c.169del NP_001074470.1:p.Tyr57Metfs*23

Allele 2

exon2 | exon3

agatgcagcacagcagatttaggaccgataagcctcaattggtttgaggagctttcctca
R C S T A D L G P I S L N W F E E L S S
gaagccccccatacaattctgaacctccggaggaatctgagtataagccccacggt-at
E A P P Y N S E P P E E S E Y K P H G M
gaaccacagctgtttaaaacaccacagaggaatccccctaccatcagtttgcttcaact
N H S C L K H H R G I P P T I S L L Q L
ccaataatgttcaaa
Q *

b

Brca2 wt
reference

NM_001081001.2:c.=, NP_001074470.1:p.=

exon10 | exon11 exon12

aatgtaaattcaggtataccagattcttct.........gcagttggacaacccccaatcaaaaga
N V N S G I P D S S A V G Q P P I K R

HGS, HGS2-OR
*Brca2del*

NM_001081001.2:c.1877_6685del, NP_001074470.1:p.Gly626_Val2228del

exon10 | exon12

aatgtaaattcaggacaacccccaatcaaaaga
N V N S G Q P P I K R

**Supplementary Figure 3**

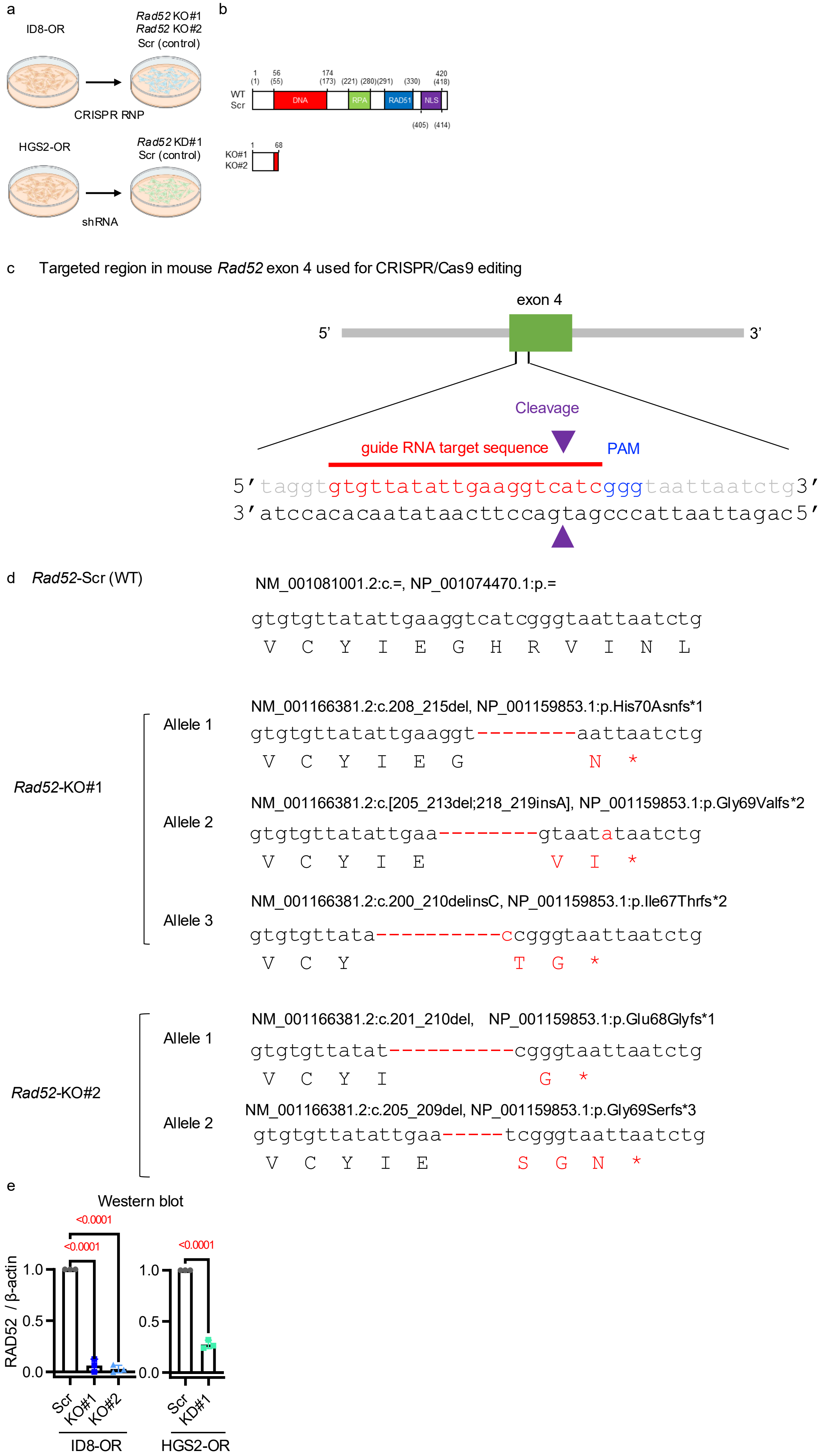

Supplementary Figure 4

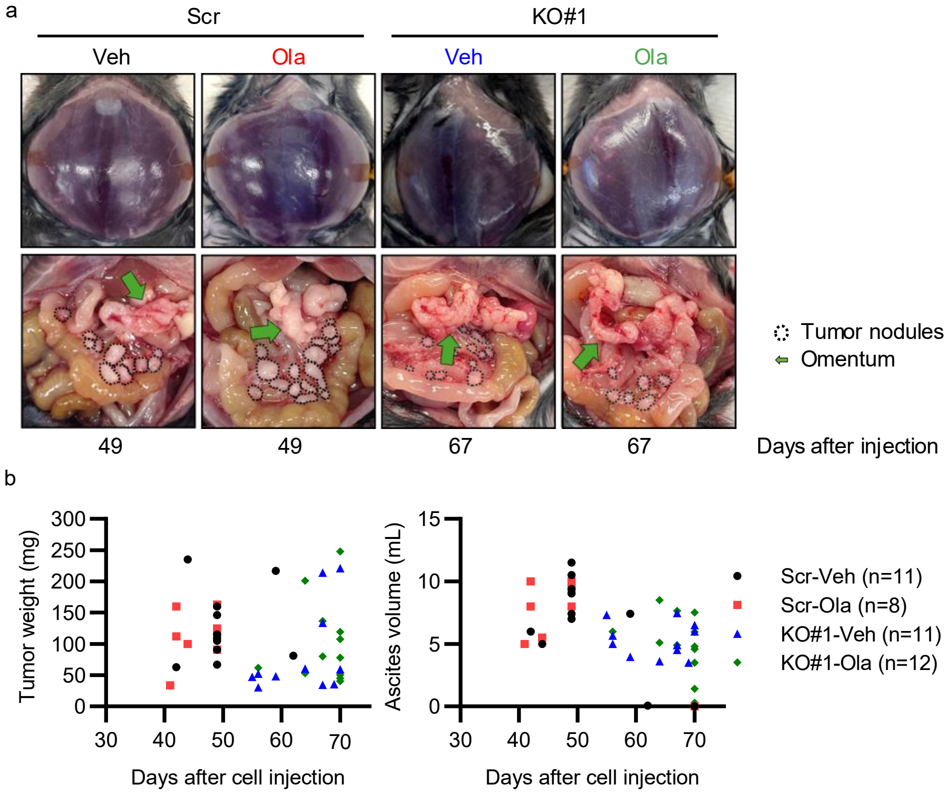

**Supplementary Figure 5**

a

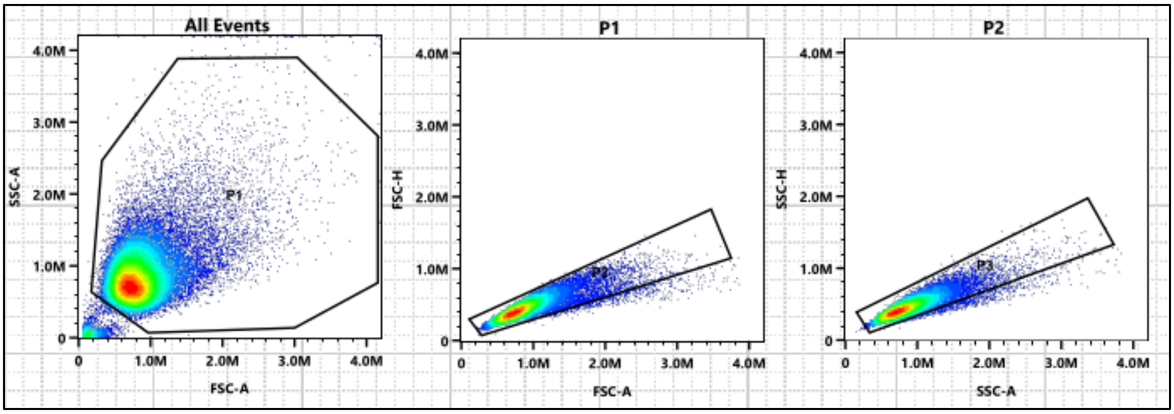

b SSA reporter assay. hprtSAGFP(+), pCBASceI (+). ID8-OR, ID8-PB, ID8-P

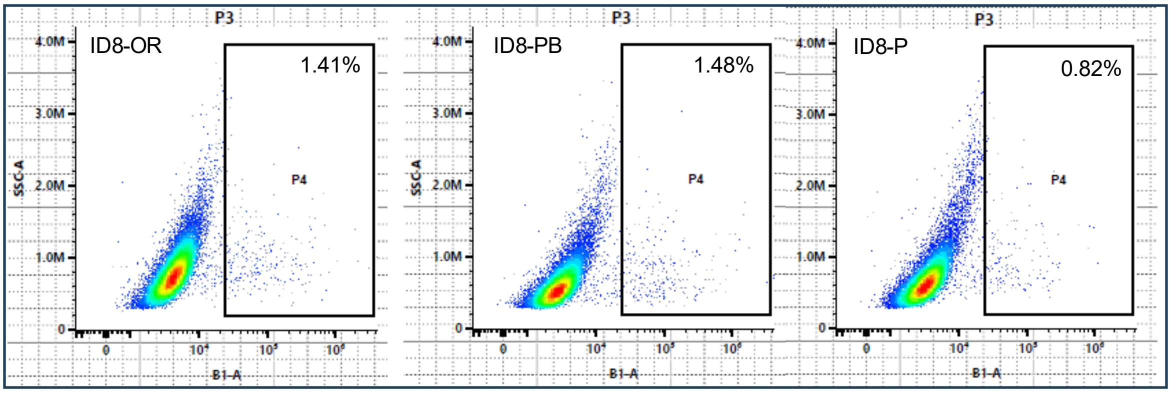


hprtSAGFP(+), pCBASceI (-). ID8-OR, ID8-PB, ID8-P

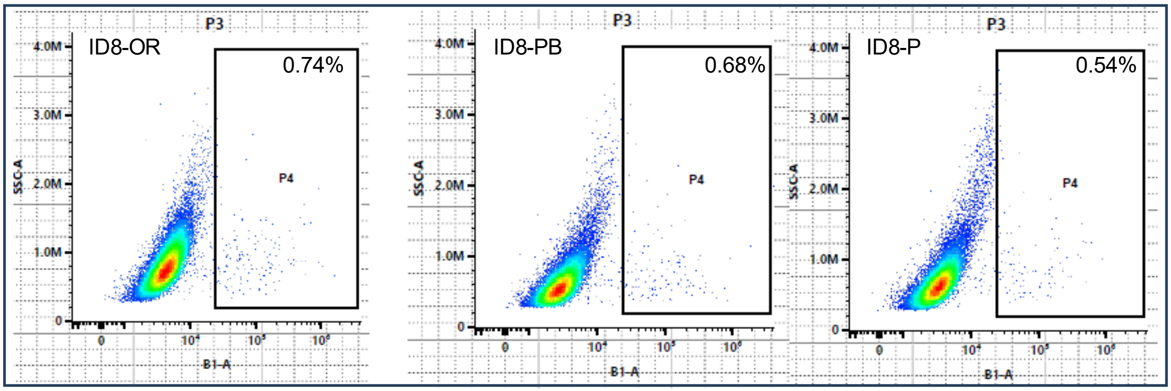


c SSA reporter assay. hprtSAGFP(+), pCBASceI (+). HGS2-OR, HGS2

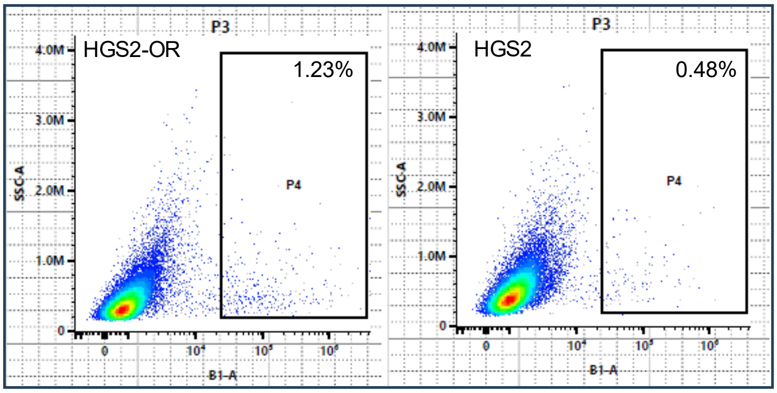


hprtSAGFP(+), pCBASceI (-). HGS2-OR, HGS2

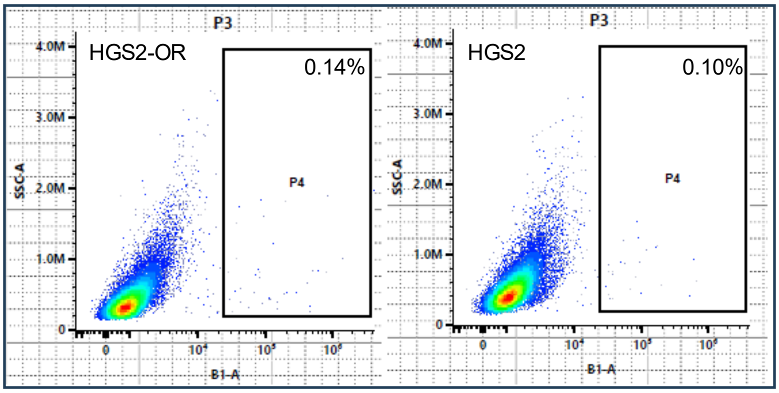

d SSA reporter assay. hprtSAGFP(+), pCBASceI (+). ID8-Scr, ID8-KO#1, ID8-KO#2

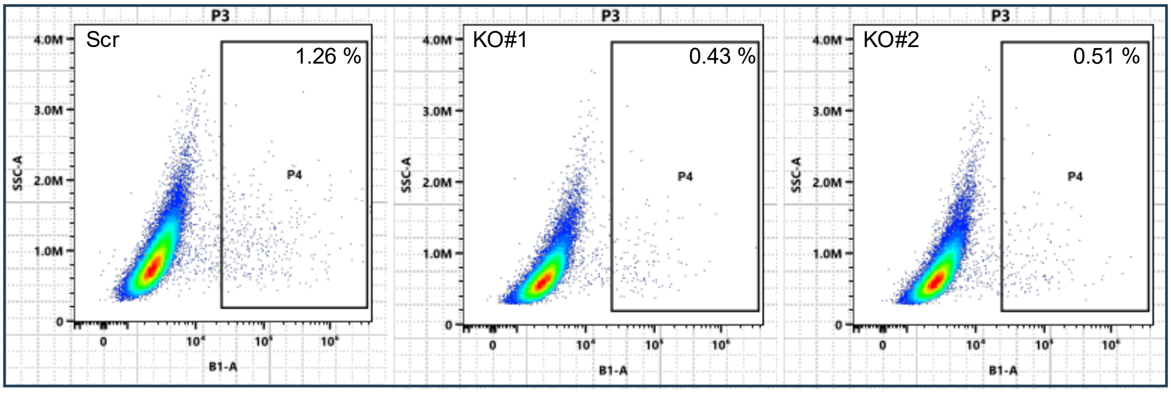


hprtSAGFP(+), pCBASceI (-). ID8-Scr, ID8-KO#1, ID8-KO#2

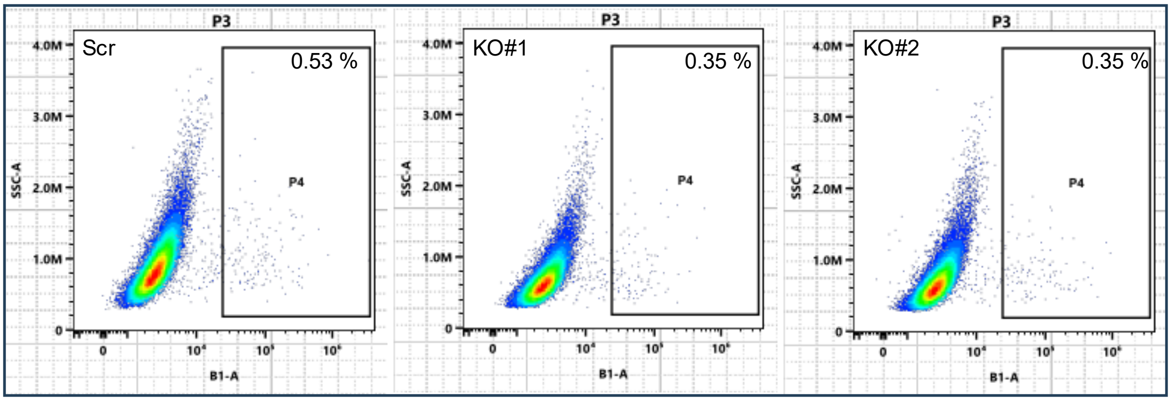


e SSA reporter assay. hprtSAGFP(+), pCBASceI (+). HGS2-Scr, HGS2-KD#1

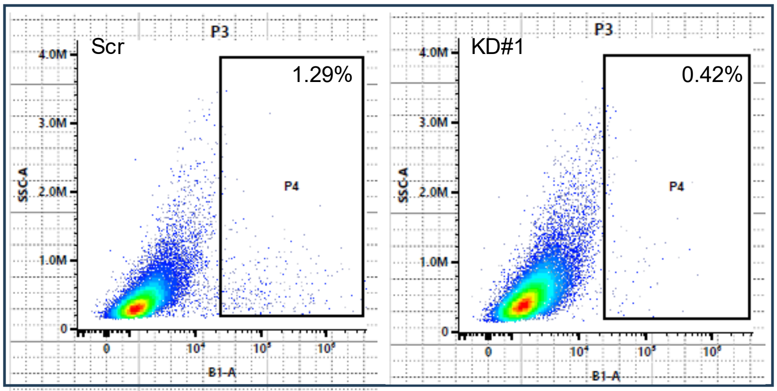


hprtSAGFP(+), pCBASceI (-). HGS2-Scr, HGS2-KD#1

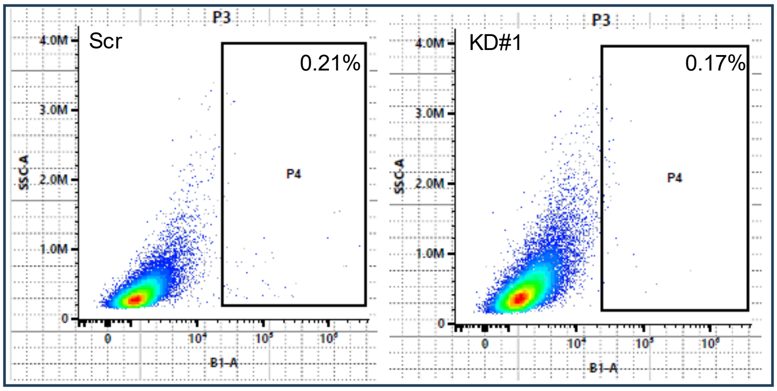

f HR reporter assay. pDR-GFP(+), pCBASceI (+). ID8-OR, ID8-PB, ID8-P

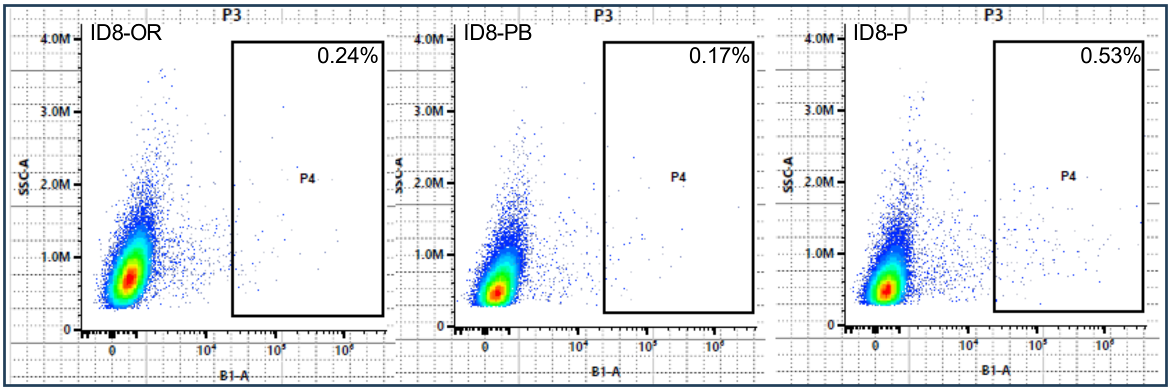


pDR-GFP(+), pCBASceI (-). ID8-OR, ID8-PB, ID8-P

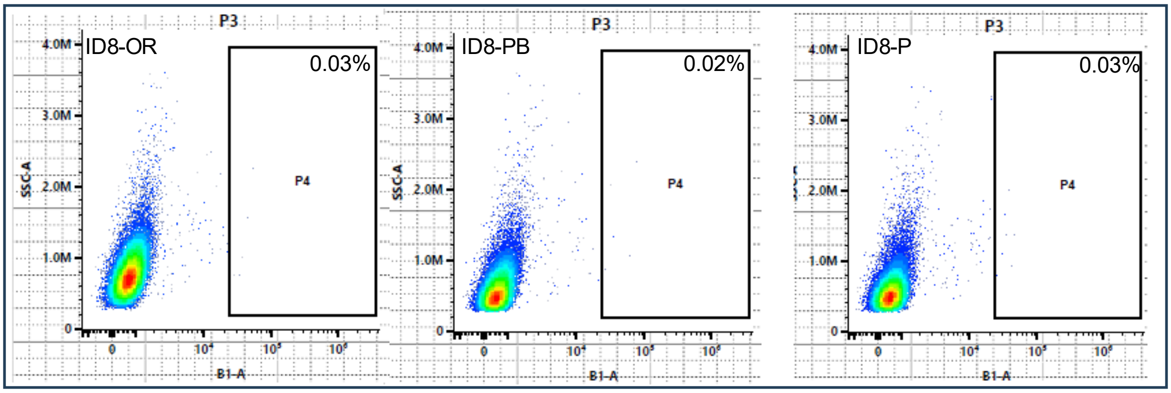


g HR reporter assay. pDR-GFP(+), pCBASceI (+). HGS2-OR, HGS2

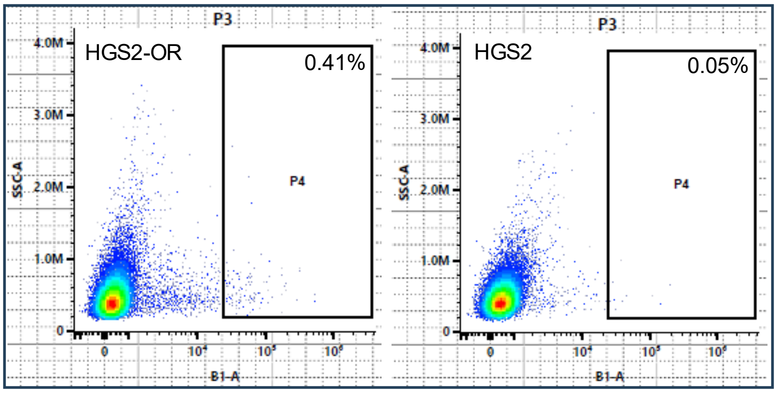


pDR-GFP(+), pCBASceI (+). HGS2-OR, HGS2

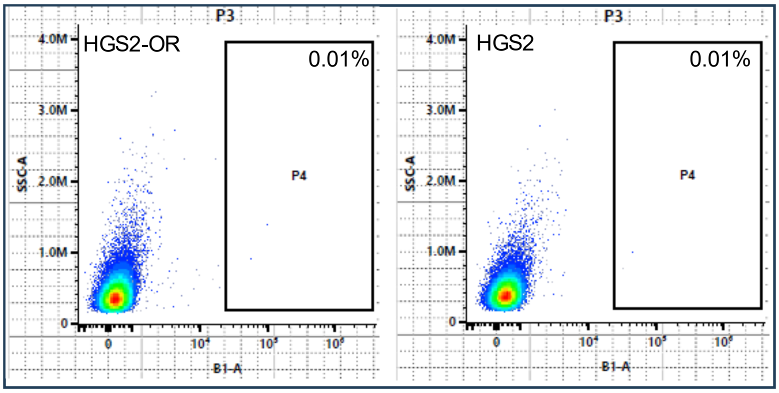

h HR reporter assay. pDR-GFP(+), pCBASceI (+). ID8-Scr, ID8-KO#1, ID8-KO#2

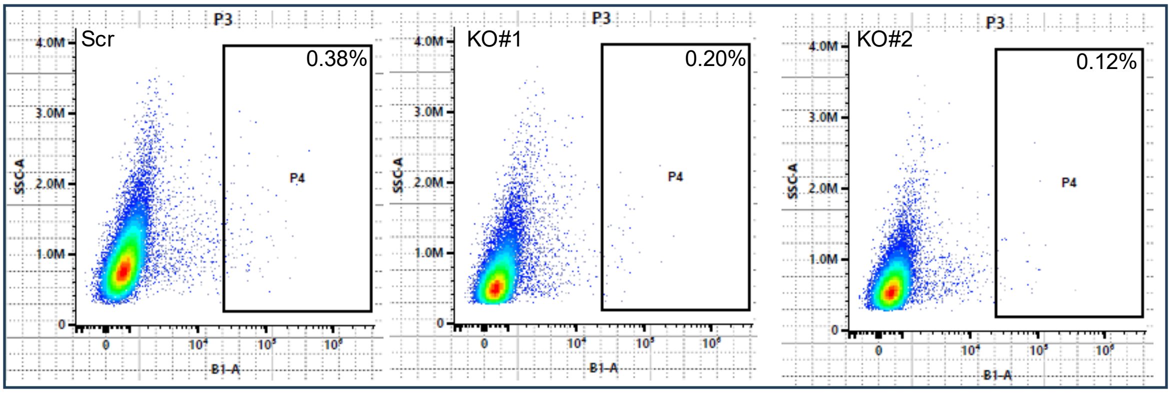


pDR-GFP(+), pCBASceI (-). ID8-Scr, ID8-KO#1, ID8-KO#2

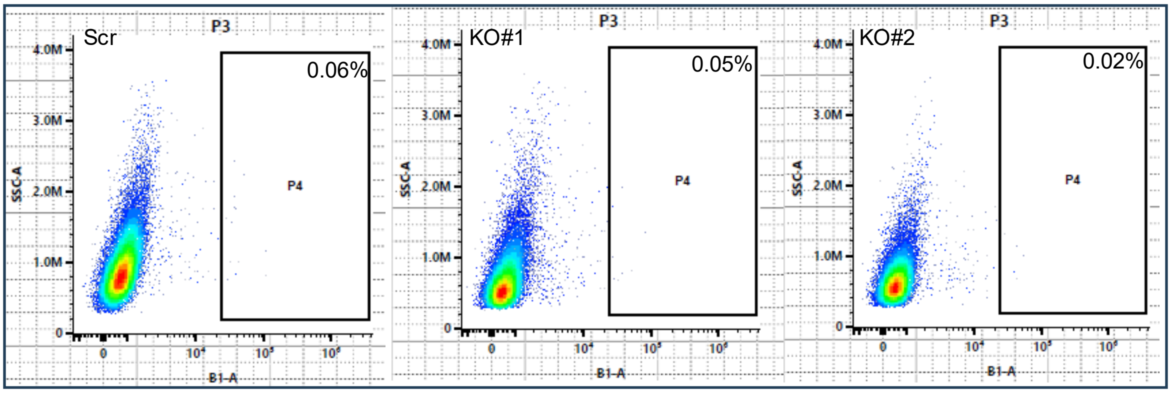


i HR reporter assay. pDR-GFP(+), pCBASceI (+). HGS2-Scr, HGS2-KD#1

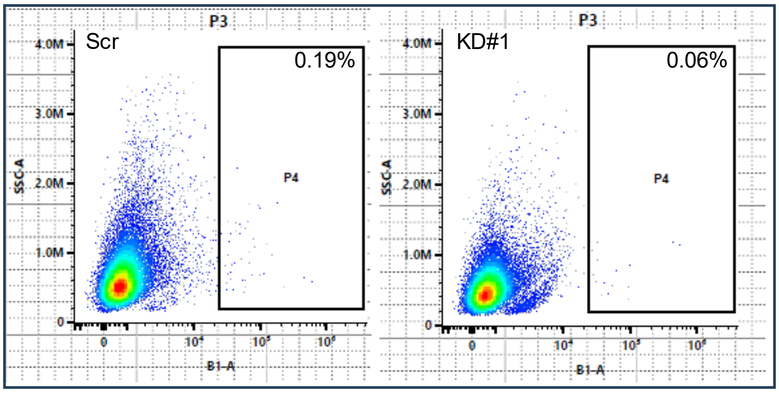


pDR-GFP(+), pCBASceI (-). HGS2-Scr, HGS2-KD#1

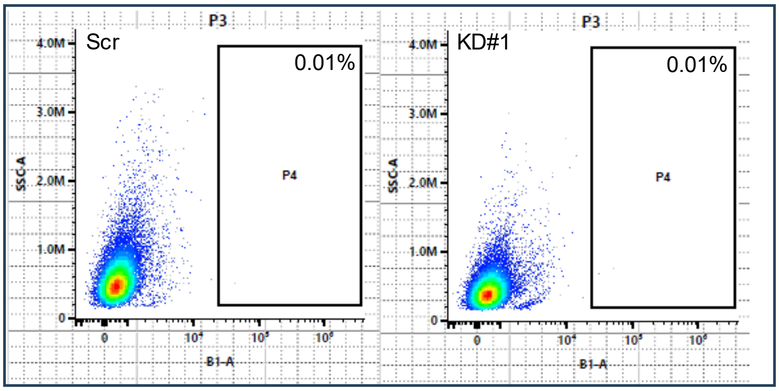

j Untransfected. ID8-P, ID8-PB, ID8-OR

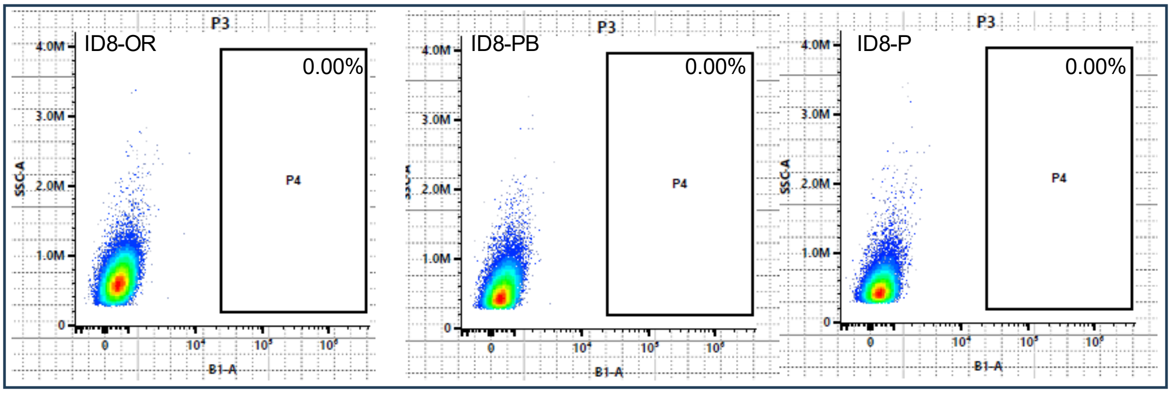


Untransfected. HGS2, HGS2-OR

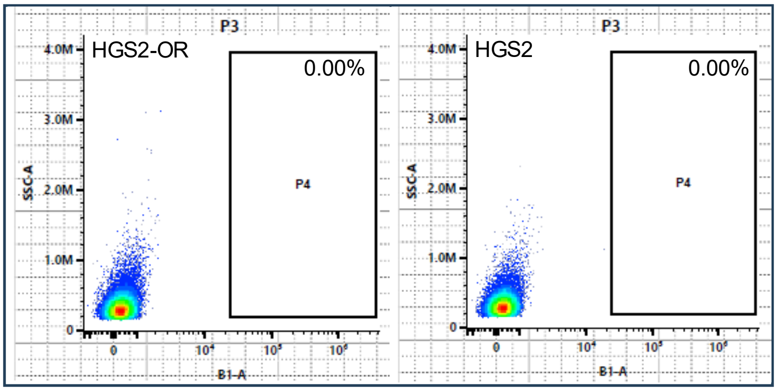


Untransfected. ID8-Scr, ID8-KO#1, ID8-KO#2

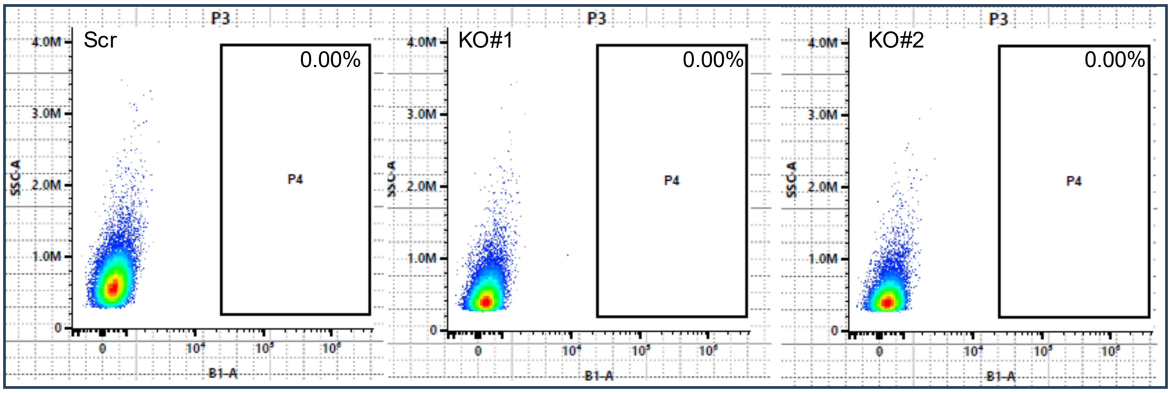


Untransfected. HGS2-Scr, HGS2-KD#1

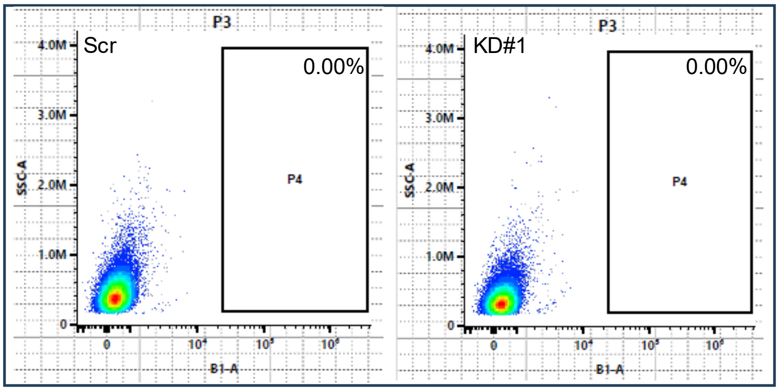

**Supplementary Figure 6**

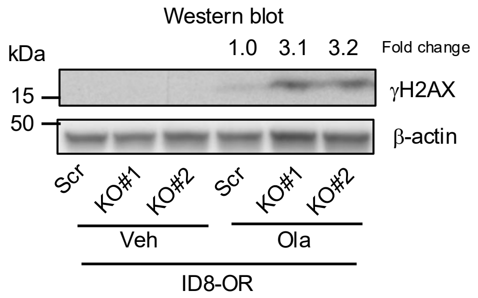

### Supplementary Figure 7

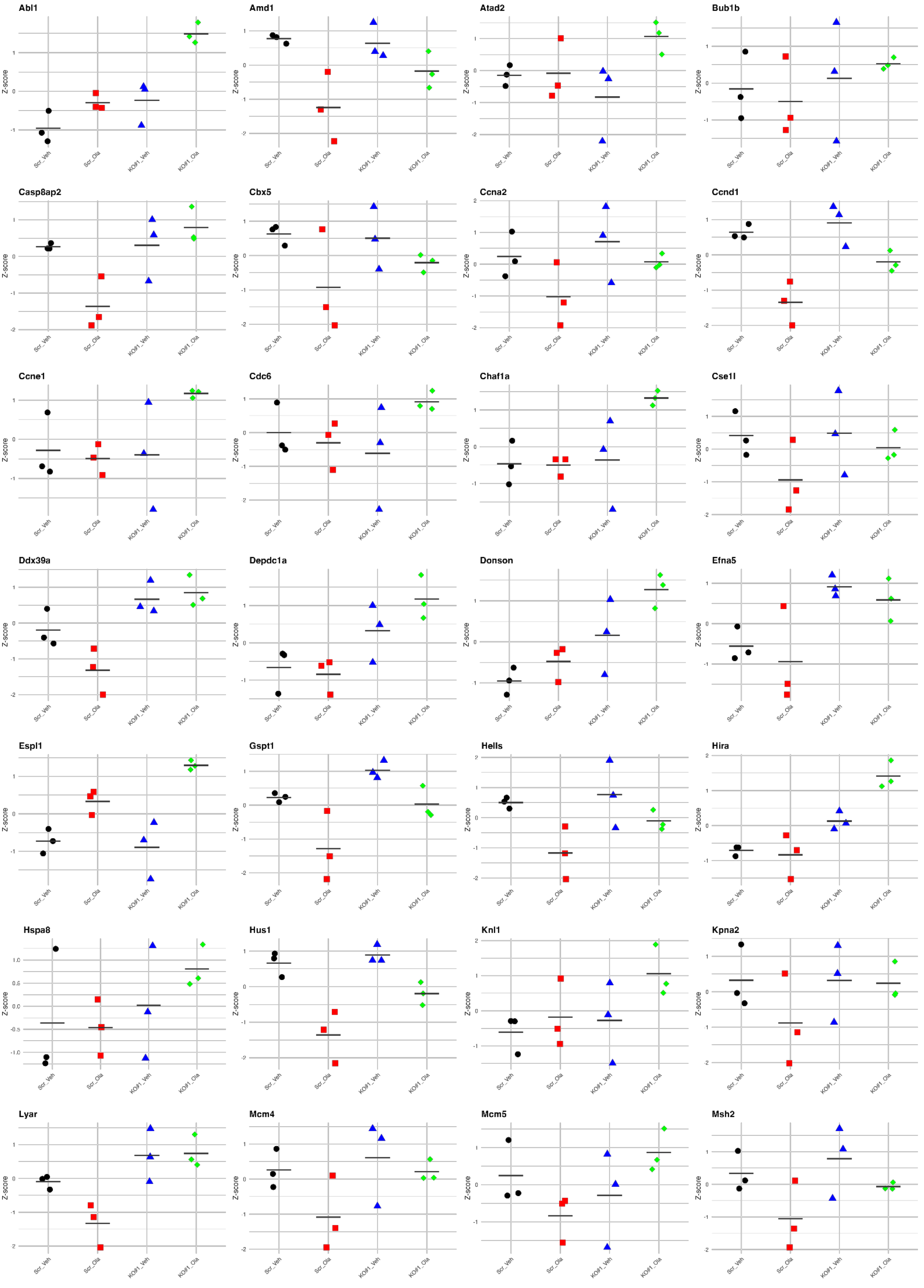

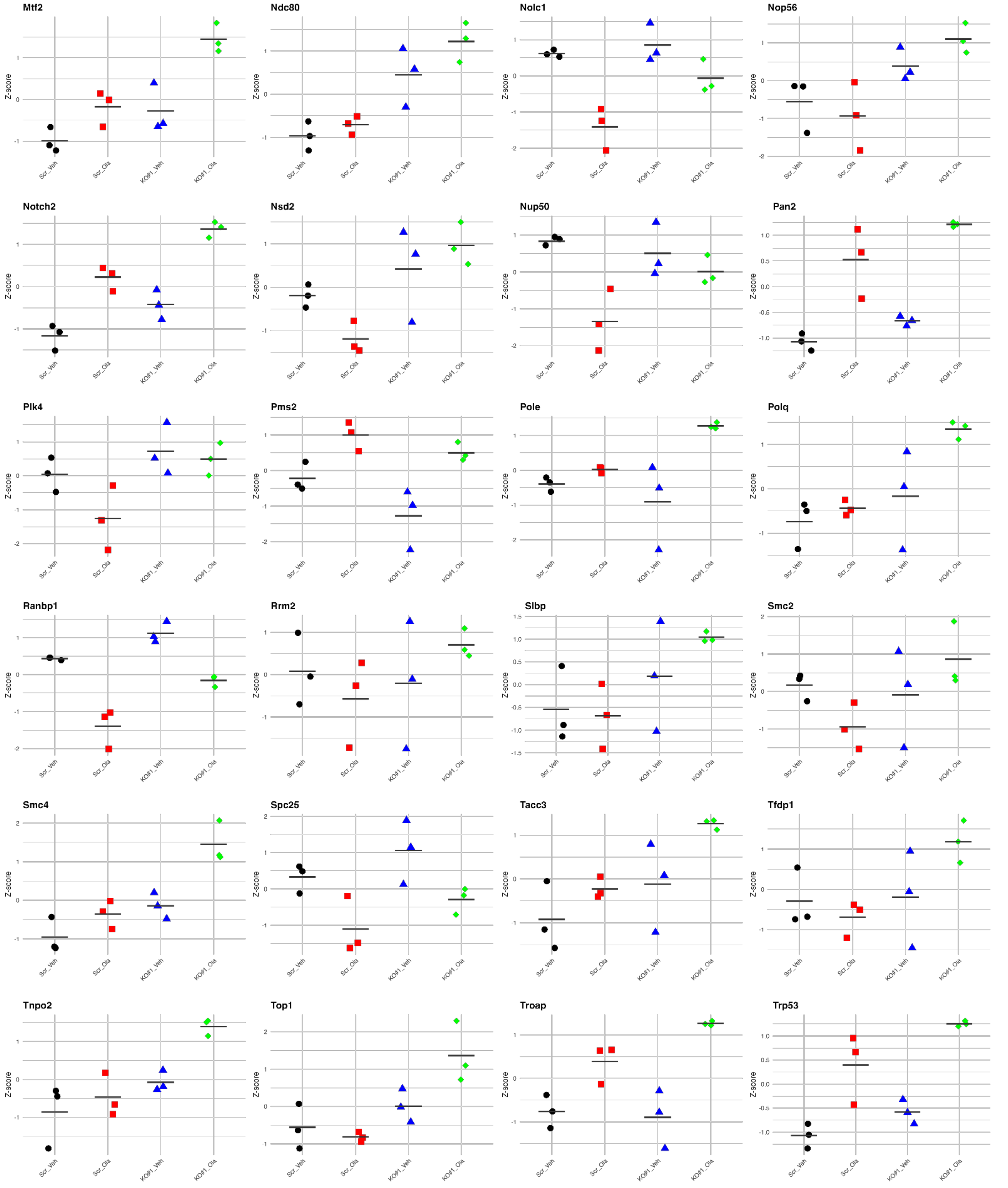
Mtf2
Ndc80
Nolc1
Nop56
Notch2
Nsd2
Nup50
Pan2
Plk4
Pms2
Pole
Polq
Ranbp1
Rrm2
Slbp
Smc2
Smc4
Spc25
Tacc3
Tfdp1
Tnpo2
Top1
Troap
Trp53
Z-score
Scr_Veh
Scr_Ola
KO#1_Veh
KO#1_Ola

**Supplementary Figure 8**

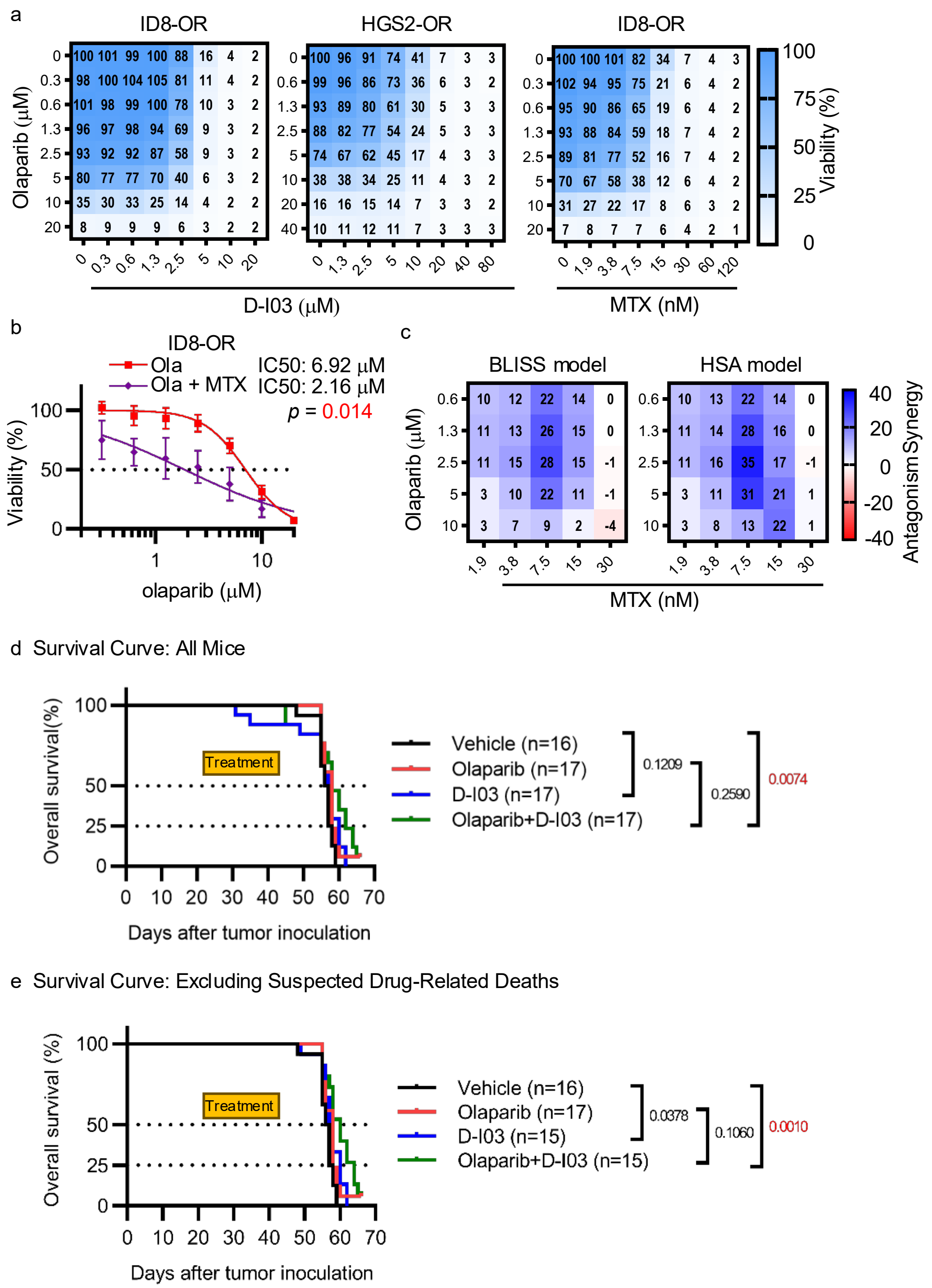

Supplementary Table 1: Patient characteristics of TMA cohort.

| | **Total (n=72)** | **RAD52 low (n=36)** | **RAD52 high (n=36)** | **P-value** |
|---|---|---|---|---|
| Age (years) | 59.6 ± 10.2 | 59.9 ± 9.8 | 59.4 ± 10.8 | 0.83 |
| FIGO stage | | | | 0.20 |
| I-II | 3 (4.2) | 3 (8.3) | 0 (0) | |
| III-IV | 64 (88.9) | 30 (83.3) | 34 (94.4) | |
| Missing | 5 (6.9) | 3 (8.3) | 2 (5.6) | |
| *BRCA* Mutation | | | | 0.30 |
| None | 32 (44.4) | 13 (36.1) | 19 (52.8) | |
| *BRCA* | 19 (26.4) | 12 (33.3) | 7 (19.4) | |
| *BRCA1* | 10 | 6 | 4 | |
| *BRCA2* | 2 | 2 | 0 | |
| *BRCA1+2* | 1 | 0 | 1 | |
| No data | 6 | 4 | 2 | |
| No testing/Missing | 21 (29.2) | 11 (30.6) | 10 (27.8) | |
| Neoadjuvant Chemotherapy | | | | |
| No | 72 (100) | 36 (100) | 36 (100) | |
| Cytoreduction Status | | | | >0.99 |
| Optimal | 55 (76.4) | 28 (77.8) | 27 (75.0) | |
| Suboptimal | 13 (18.1) | 6 (16.7) | 7 (19.4) | |
| Missing | 4 (5.6) | 2 (5.6) | 2 (5.6) | |
| PARP inhibitor Use | | | | 0.31 |
| YES | 22 (30.6) | 13 (36.1) | 9 (25.0) | |
| NO | 49 (68.1) | 22 (61.1) | 27 (75.0) | |
| Missing | 1 (1.4) | 1 (2.8) | 0 (0) | |
| Median Follow-up Months (IQR) | 64.0 (37.8–89.8) | 68.0 (50.3–89.8) | 59.0 (25.3.3–90.5) | 0.22 |

Data are n (%) unless stated otherwise. ± Denotes standard deviation Statistical analyses were performed as follows: age was compared using the Student's t-test; follow-up duration was analyzed using the Mann–Whitney U test; and categorical variables were compared using Fisher's exact test. IQR, Interquartile Range.

Supplementary Table 2 Cell lines used in this study.

| Cell line | Lineage | Genetic background | Notes | Ref. |
|---|---|---|---|---|
| ID8-P | Mouse | *Trp53*$^{-/-}$ | | 82 |
| ID8-PB | Mouse | *Trp53*$^{-/-}$, *Brca2*$^{-/-}$ | Derived from ID8-P | 82 |
| ID8-OR | Mouse | *Trp53*$^{-/-}$, *Brca2*$^{-/-}$ | Olaparib-resistant. Derived from ID8-PB. | 48 |
| ID8-Scr | Mouse | *Trp53*$^{-/-}$, *Brca2*$^{-/-}$ | Scrambled gRNA control. Derived from ID8-PB-OR | |
| ID8-*Rad52*KO#1, *Rad52*KO#2 | Mouse | *Trp53*$^{-/-}$, *Brca2*$^{-/-}$, *Rad52*$^{KO}$ | Rad52-knockout clone #1 and #2 via CRISPR/Cas9. Derived from ID8-PB-OR | |
| HGS2 | Mouse | *Trp53*$^{-/-}$, *Brca2*$^{-/-}$, *Pten*$^{-/-}$ | | 83 |
| HGS2-OR | Mouse | *Trp53*$^{-/-}$, *Brca2*$^{-/-}$, *Pten* $^{-/-}$ | Olaparib-resistant. Derived from HGS2 | 49 |
| HGS2-Scr | Mouse | *Trp53*$^{-/-}$, *Brca2*$^{-/-}$, *Pten* $^{-/-}$ | Non-targeting shRNA control. Derived from HGS2-OR | |
| HGS2-*Rad52*KD#1 | Mouse | *Trp53*$^{-/-}$, *Brca2*$^{-/-}$, *Pten* $^{-/-}$ | Rad52-knockdown clone #1. Derived from HGS2-OR | |
